# Chromatin accessibility differences between the hybrids of *nasuta-albomicans* complex of *Drosophila*

**DOI:** 10.1101/2025.06.26.661782

**Authors:** Radhika Padma, Ajinkya Bharatraj Patil, Nagarjun Vijay

## Abstract

Understanding how chromosomal rearrangements (CRs) interact with epigenetic changes to drive speciation is a fundamental question in evolutionary biology. CRs are key contributors to chromosome evolution and can play a pivotal role in reproductive isolation. The *Drosophila nasuta-albomicans* species complex presents an ideal model to explore this, as *D. albomicans* possesses neo-sex chromosomes formed by Robertsonian fusion of chr3L, chr3R, and a sex chromosome, unlike its sister species *D. nasuta*. In this study, we investigate CRs influence on accessible chromatin (AC) and its relationship with genetic differentiation. We used ATAC-seq to decipher ACs of testis in two hybrids of *D. albomicans* and *D. nasuta*, and genome-wide fixation index (F_ST_) scans of *D. albomicans* and *D. nasuta* to identify regions of genetic differentiation. Our analyses revealed that chromosome 4 (Muller F) harbors the largest number of differentially accessible regions (∼97 Kb), which coincide with peaks in F_ST_. Moreover, changes in ACs were associated with differential transcription factor (TF) binding across the genome. These results suggest that CRs can drive epigenomic divergence in hybrids, particularly on Muller F, and chromatin-level changes may play a key role in reproductive isolation. Our study provides an example of how chromosomal and epigenetic architecture interact in the early stages of speciation.

## Introduction

Genome regulation, a curator of the genome, uses multiple means to initiate and sustain tissue-specific functionalities in a multicellular organism, thus playing a pivotal role in evolutionary diversification (Wray 2003). For example, beak morphological differences seen in Darwin’s Finches (Abzhanov et al. 2004); pelvic reduction in sticklebacks (Chan et al. 2010); wing patterns in dipteran insects (Prud’homme et al. 2007); regulatory divergence within and between species of Drosophila (Wittkopp et al. 2008). Incidentally, epigenetic components govern genome regulation: DNA methylation, histone tail modifications, chromatin accessibility, and DNA architecture (Holtzman and Gersbach 2018). We are primarily interested in chromatin accessibility, a dynamic property that determines and sustains cellular identity (Klemm et al. 2019). For instance, accessible chromatin (AC) allows transcriptional and regulatory machinery to interact with the DNA physically, and they, in turn, are regulated by the nucleosomes (Allis and Jenuwein 2016).

Nucleosome distribution across the genome is an integral part of genome regulation, determining cells’ developmental trajectory and tissue-specific identity. Thus, nucleosome density and depletion across the genome represent inactive heterochromatin (Sun et al. 2001) and active euchromatin (Lee et al. 2004). Not all the heterochromatin regions found in the genome are perennial; based on the nature of inactivation (Richards and Elgin 2002), they can be (a) constitutive, where the inactivation is permanent and consistent across cell types, and (b) facultative, when the inactivation is transient and a hallmark of cell-type and locus-specific, for example, X-chromosome inactivation (Allshire and Madhani 2018). The constitutive heterochromatin guards the genome against deleterious effects of rearrangement, recombination, and transposons (Janssen et al. 2018). They can breakdown due to chromosomal rearrangement (CR), leading to position effect variegation (PEV); a phenomenon first identified in Drosophila, later in others (Muller 1930; McClintock 1953), which modifies the gene expression of organisms (Elgin and Reuter 2013; Harewood and Fraser 2014). Even though heterochromatin is inaccessible to the transcription factors (TFs), a class of TFs known as pioneer TFs bind to them and change the accessibility (Balsalobre and Drouin 2022; Duan et al. 2021).

CRs as an influential evolutionary player, reshuffle and disrupt gene order/synteny (Bhutkar et al. 2008), resulting in changing the gene expression (Harewood and Fraser 2014; Stewart and Rogers 2019), regulation (Goettel and Messing 2010), phenotype (Muller 1930), adaptation (Guerrero and Kirkpatrick 2014), and speciation (Rieseberg 2001; Ayala and Coluzzi 2005; Kirkpatrick and Barton 2006). Numerous events of CR in insects hint at their possible role in the diversification of insects (de Vos et al. 2020; Lukhtanov et al. 2011; Mathers et al. 2021; Bhutkar et al. 2008; Ranganath and Hfigele 1982). The role of CR in promoting genetic variation and differentiation is complex (Navarro and Barton 2003; Ayala and Coluzzi 2005). Therefore, understanding the evolution of interactions between CRs, chromatin organization, and genome regulation is an important question.

Affordable DNA sequencing technologies allowed large-scale investigation of various aspects of biology, for example, how genetic changes accumulate between species (Holsinger and Weir 2009; Corbett-Detig and Hartl 2012; Poelstra et al. 2014; Kou et al. 2020; Hollox et al. 2022). These studies identified high levels of genetic differentiation in short genomic regions commonly called speciation islands (Turner et al. 2005). These speciation islands are interpreted either as barriers to gene flow (Feder et al. 2012; Poelstra et al. 2014) or due to varying degrees of linked selection (Comeron et al. 2012; Burri et al. 2015; Wolf and Ellegren 2017). In contrast to these widely studied genetic differentiation islands, no study has demonstrated an extreme accumulation of between-species epigenetic changes in genomic islands, to the best of our knowledge. Hence, it is unknown whether such epigenetic islands of speciation exist.

We are interested in effects of CR event seen in *Drosophila albomicans* in hybrids of *D. albomicans* and *D. nasuta* (**Fig. 1A**).Where, *D. albomicans* has a centric fusion of three Muller Elements (CF-3M) leads to forming a neo-sex-chromosome (Ranganath and Hfigele 1982). On the other hand, *D. nasuta,* a closely related species of *D. albomicans,* does not display this CF-3M CR, making them an ideal system for understanding the changes driven by a CR in hybrids. Both species belong to the *D. immigrans* species group in the subgenus *Drosophila.* The *D. immigrans* species group has experienced a burst of speciation events during a short time, resulting in species with little to no morphological differences and reproductive isolation (Wilson et al. 1969; Kitagawa et al. 1982). In recent years, *D. albomicans* have garnered ample attention leading to the exploration of the genome (Zhou et al. 2012), expression profile across tissue types (Gibilisco et al. 2016; Nozawa et al. 2021), male achiasmy (Satomura and Tamura 2016; Wei and Bachtrog 2019), inversions (Mai and Bachtrog 2021), and transposon density (Wei et al. 2021). However, the effect of centric fusion on regulome and its significance on speciation is understudied. Similarly, the association of speciation islands with regulation is not known yet. With this study, we hope to understand the impact of CRs on the genome and how it changes the chromatin landscape and regulation in hybrids.

**Figure 1:**
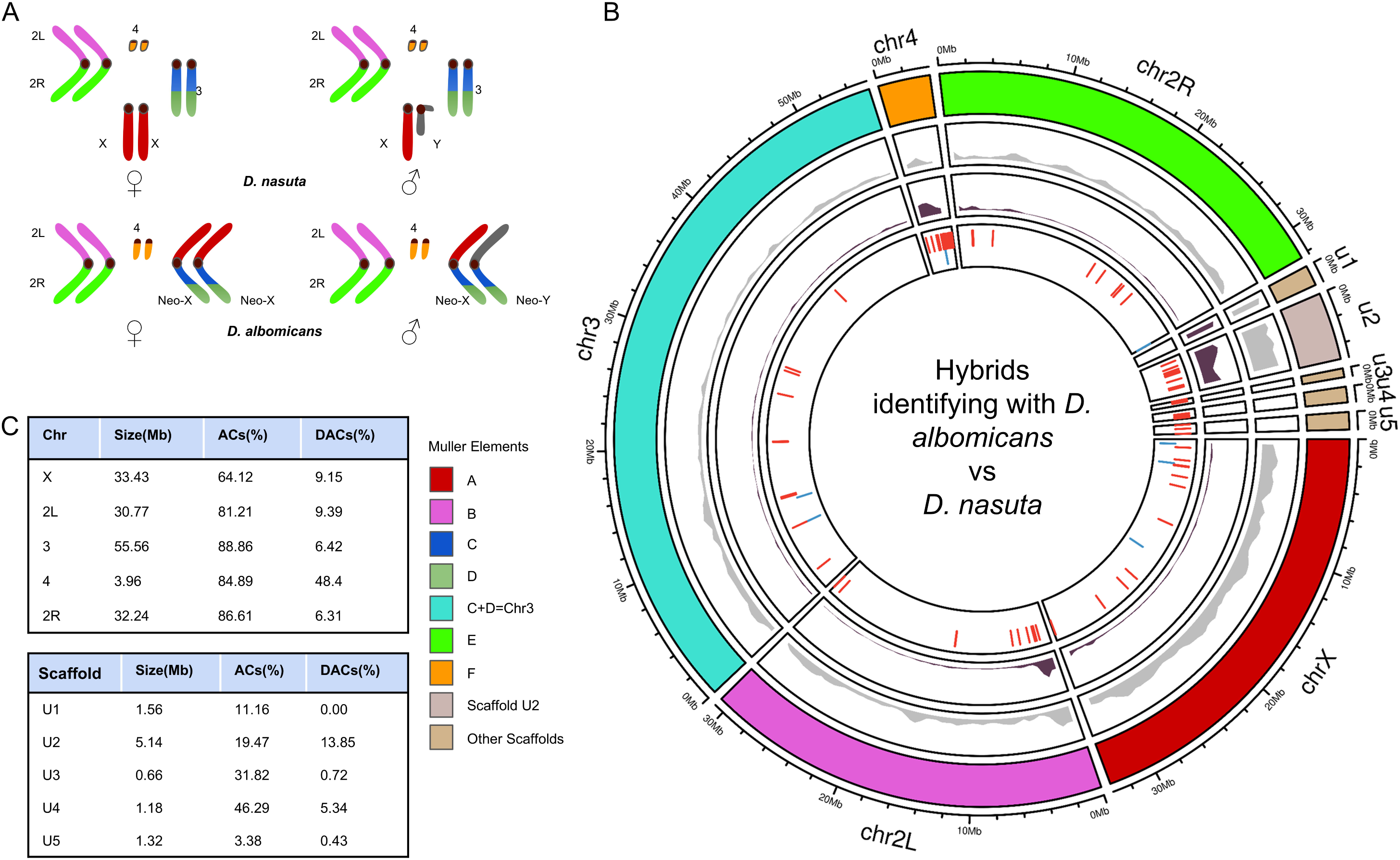
Karyotype and testes’ chromatin landscape of *D. albomicans* and *D. nasuta*. **A** Karyotype of male and female *D. albomicans* and *D. nasuta.* The karyotype of *D. nasuta* shows ancestral chromosome complement, and *D.albomicans* shows rearranged chromosomes. Where rearrangement of chromosomes leads to the formation of neo-sex-chromosomes by the fusion of Muller elements A, C, and D. The chromosome arms colored with corresponding Muller elements in both the species. **B** Accessible chromatin (ACs) of testes in hybrids identifying with *D. albomicans* and *D. nasuta,* the outermost and second outermost tracks represent chromosomes, and non-accessible chromatin density, respectively. The innermost track represents differential accessibility (DACs), blue and red bars represent high and low accessibility regions in hybrid identifying with *D. albomicans*. The second innermost track represents the transposable element density. The chromosome colors depicted Muller elements, except for chromosome 3; due to the fusion of Muller elements (C and D) and unplaced scaffolds, the scaffold of the largest size (i.e., U2) represented in a different color from others. **C** Chromosome sizes with ACs and DACs by DESeq2 across *D. albomicans* genome.

To explore the accessibility variation between the hybrids of *D. albomicans* and *D. nasuta*, we have used the ATAC-seq of adult testes with biological replicates. The genome-wide ATAC-seq landscape revealed the presence of accessibility variation across chromosomes and implicated Muller F as one of the prime candidates for an epigenetic speciation island. Further population genetic study of public datasets found this epigenetic island overlaps with islands of genetic differentiation. Additionally, the prediction of transcription factor binding sites (TFBS) suggests the accessibility difference between the two hybrids. Our findings determine that genetic differentiation, accessibility differences, and TFBS variation are interlinked.

## Materials and Methods

### ATAC-seq data generation

The hybrids identifying with *D. albomicans* and *D. nasuta* (see **Supplementary Text** for details) were collected from the Drosophila Stock Center, University of Mysore, India, and maintained on a standard culture medium at 25°C. The testis of two species was harvested from 5-10 days old males with PBS (pH 7.4) and stored in Recovery™ Cell Culture Freezing Medium (cat no 12648010, Gibco). The study consists of four samples, replicates from each species, and each sample consists of 25-30 individual testis. We used Active motif’s ATAC-seq kit (cat no. 53150) to create ATAC-seq libraries of PE-150 with 450±50 bp insert size sequenced on the Illumina NovaSeq 6000. The sequenced data is available at ENA under the project PRJEB46771 (**Supplemental Table S1**).

### Data analysis

Our analysis involves three steps (a) **pre-core analysis**, (b) **core analysis**, and (c) **advanced analysis** (**Supplemental Fig. 1**), and the analysis pipeline is available on GitHub (https://github.com/RadPa/ATAC-seq.git).

### Pre-core analysis

The raw reads passed through customary FastQC v0.11.9 (Andrews and others 2010), which reported the presence of adapters. Thus, Cutadapt v2.8 (Martin 2011) trimmed these adapters, and MultiQC v1.11 (Ewels et al. 2016) aggregated the fastqc reports. The adapter pruned reads aligned to the *D. albomicans* (RefSeq: GCF_009650485.1_drosAlbom15112) genome using Bowtie v2-2.4.2 (Langmead et al. 2009) with the following parameters -X 750 -very-sensitive. After read alignment, Samtools v1.13 (Li et al. 2009) removed mitochondrial reads and MarkDuplicates of GATK v4.2.2.0 (Van der Auwera and O’Connor 2020) marked the duplicates. After the initial quality checks, 80% of reads mapped to the genome of *D. albomicans*, followed by various filtering (**Supplemental Table S1**). We assessed the library quality and fragment length distribution of aligned reads using the ATACseqQC (Ou et al. 2018), an R package (**Supplemental Fig. 2 and 3**). Additionally, ATACseqQC shifted the aligned reads, i.e., + strand reads offset by +4 bp and – strand reads offset by −5 bp.

### Blacklist

ENCODE (Amemiya et al. 2019) suggests the removal of unstructured high signal regions of the genome called blacklist for functional genomic analysis before peak calling. We used public DNA-seq datasets for *D. albomicans* and *D. nasuta* (**Supplemental Table S2**) to establish a blacklist for the species. The datasets were mapped to *D. albomicans* genome (RefSeq: GCF_009650485.1_drosAlbom15112) using bwa-mem v0.7.17 (Li and Durbin 2009), Samtools v1.13 used for post-processing of aligned reads, and MarkDuplicates of GATK v4.2.2.0 (Van der Auwera and O’Connor 2020) removed the optical duplicates. The umap v1.1.1 tool (Karimzadeh et al. 2018) (https://github.com/hoffmangroup/umap) identifies uniquely mappable regions for the genome, and Blacklist (https://github.com/Boyle-Lab/Blacklist) utilizes umap regions to build blacklist regions. For our analysis, we used merged blacklist regions of *D. albomicans* and *D. nasuta*, using bedtools v2.30.0 (Quinlan and Hall 2010) to represent both species. We estimated 4.9% of the genome as blacklist and showed the importance of using more datasets to detect such regions (**Supplemental Table S3**).

### Core Analysis

#### Peak calling

We have used HMMRATAC v1.2.10 (Tarbell and Liu 2019) to identify the accessible chromatin regions of testis in our datasets. The program uses the reads and genome as requirements; we have used the shifted reads from ATACseqQC, *D. albomicans* chromosome size, and blacklist, as described above. Most of the parameters were unchanged except for minimum mapping quality scores, changed to 20; fold change ranges, -u and -l (**Supplemental Table S4**). We selected the sensitive models producing the highest number of peaks per dataset for the downstream analysis (**Supplemental Fig. 4**). The peak files were filtered for nucleosome-free regions, as HMMRATAC identifies both nucleosome-enriched and nucleosome-free regions in the datasets; based on a minimum of 10 read support.

### Differential Accessibility between two hybrids

To find the presence of differential accessibility, we concatenated and sorted peak files from four datasets by Unix utility and merged using bedtools. Then bedtools multicov command generated the count matrix for all the ACs across datasets; the count matrix filtered for regions less than 10 reads support across four datasets. DESeq2 (Love et al. 2014) and edgeR (Robinson et al. 2010) used the count-matrix to identify differential accessible regions between two hybrids. The DACs from the analysis intersected with gene annotations that were enhanced with AGAT v0.8.0 (Dainat et al. 2021) (https://github.com/NBISweden/AGAT) and sorted by gff3sort v1.0.0 (Zhu et al. 2017) (https://github.com/billzt/gff3sort).

### Advanced analysis

#### Population genetic analysis

We utilized publicly available whole-genome sequencing (WGS) datasets of various populations of *D. albomicans* and *D. nasuta* (**Supplemental Table S2**) to understand population genetic statistics. We mapped the reads to the *D. albomicans* genome (RefSeq: GCF_009650485.1_drosAlbom15112). Population-specific read groups were added to the bam alignments using AddOrReplaceReadGroups, and further sequencing duplicates were removed using MarkDuplicates of GATK v4.2.0 (Van der Auwera and O’Connor 2020). We estimated the folded site frequency spectrum (SFS) for each population using genotype likelihoods generated by ANGSD v 0.935-53-gf475f10 (Korneliussen et al. 2014). These SFS outputs of each population were used to calculate nucleotide diversity statistics using the saf2theta command of realSFS. Nucleotide diversity stats were converted into 50Kb non-overlapping window format using thetastat module of ANGSD. Two-dimensional SFS was obtained using the realSFS module of ANGSD using overlapping sites for pairwise comparisons. These 2D-SFS for each pairwise comparison between populations were used to generate per base and 50 kb non-sliding window F_ST_ metrics using the F_ST_ stats2 module of ANGSD. Generally, sex chromosomes harbor a higher differentiation than autosomes (Ellegren, H. et al. 2012). We also observed a similar pattern of higher differentiation within species pairs than between species pairs. We have excluded chrX from the analyses due to the heterogeneity in population sampling of male and female individuals.

### Transcription factor binding site analysis

To find the footprint of ATAC-seq across samples, we have used ATAC-seq specific digital genomic footprinting method, TOBIAS v0.13.2 (Bentsen et al. 2020), which discovers differential binding of transcription factor (TF) between samples. The unshifted filtered reads of replicates merged using Samtools merge, as TOBIAS does not support biological replicates. TOBIAS uses aligned reads, accessible regions identified by HMMRATAC, annotation by UROPA (Kondili et al. 2017), genome, and TF motifs from the JASPAR2022 (Castro-Mondragon et al. 2022) to find the differentially bound TF between two species. The TF network was created using TOBIAS CreateNetwork and visualized in Cytoscape 3.9.1 (Shannon et al. 2003). The network represents only 118 TFs, because of limited annotation in *D. albomicans*.

## Results

### Differentially accessible Muller F

We found 42,534 Accessible Chromatin (AC) regions across four samples, spanning 77.8% of the genome assembly with varying accessibility (**Fig. 1B**). The autosomes had a higher degree of accessibility (mean: 85.4%) than chrX (64.1%). All the autosomes had a comparable level of accessibility (see **Fig. 1C and Supplemental Table S5**). DESeq2 (149 regions) and edgeR (98 regions) identified the Differentially Accessible Chromatin (DAC) regions between hybrids of *D. albomicans* and *D. nasuta* (**Supplemental Table S6 and S7**). Both tools identified 87 common regions as DAC, which comprise 58.4% and 88.7% of all DACs determined by DESeq2 and edgeR, respectively. The remaining 41.6% and 11.2% of DACs are unique to DESeq2 and edgeR (**Supplemental Table S8**). Most of the unique DACs determined by DESeq2 are present in chr2L (25.8%) and chrX (17.7%), whereas edgeR finds unique regions in chr2L (54.5%). We observed uneven distribution of DACs across the genome, with most regions located on the chr4 (Muller F) (**Supplemental Table S5**), even though it constitutes only 2.4% of the whole genome. Muller F hosts DACs that span 97Kb of its 3.9Mb (**Fig. 2** and **Supplemental Fig. 5**), which entails 48.4% of all DAC regions identified by DESeq2 and 56.41% of edgeR (**Supplemental Table S5**). Details comparing the tools and variation of ACs and DACs across chromosomes are in **Supplemental Table S5**.

**Figure 2:**
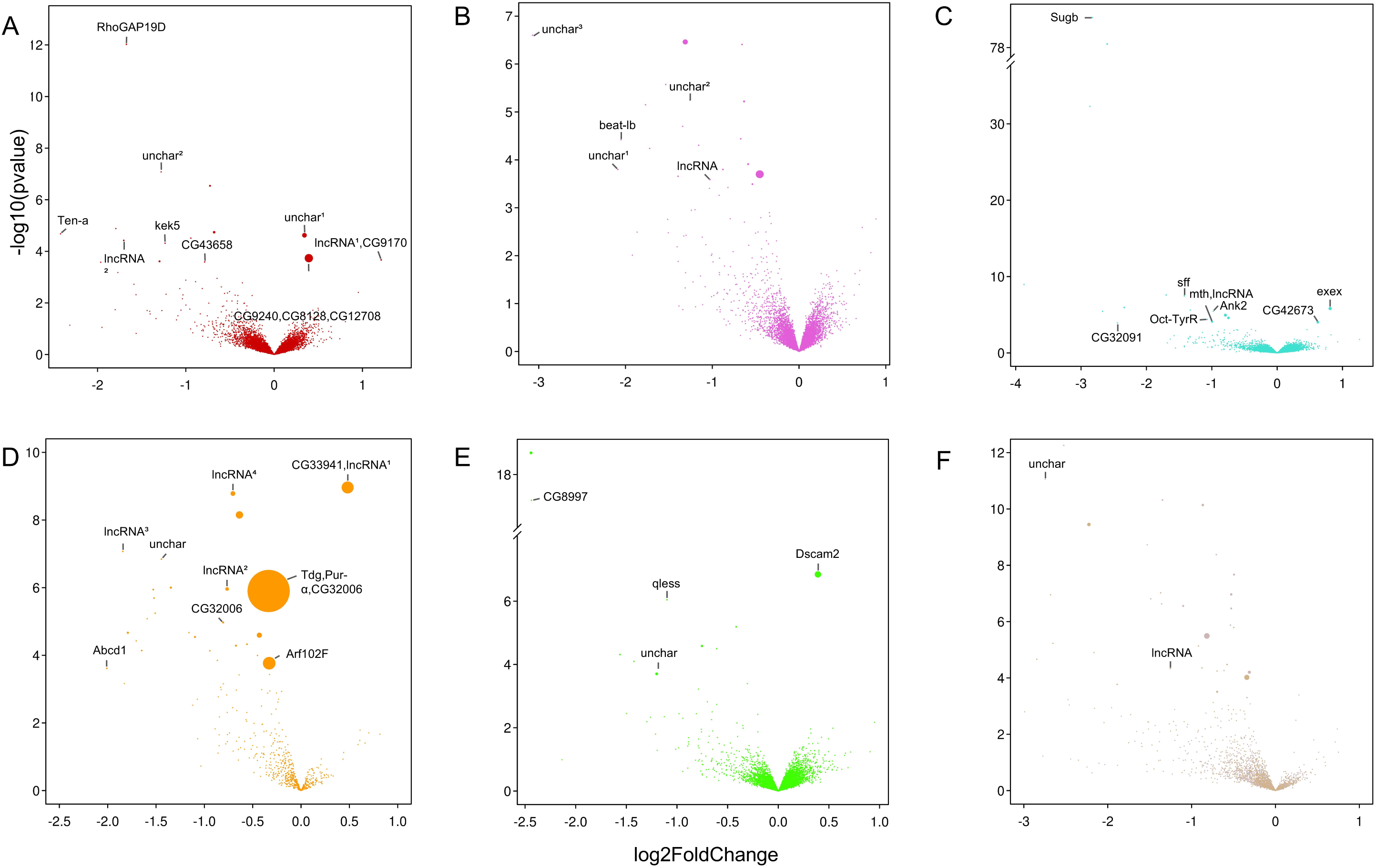
Volcano plots of differentially accessible chromatins (DACs) **A** to **F** DACs of chromosome X, 2L, 3, 4, and 2R, respectively, the x-axis denotes the log2-fold-change, while the y-axis denotes -log10 of the p-value. **C** shows the y-axis break between 38 and 76, and **E** shows the y-axis break between 8 and 16. The chromosomes are represented in Muller colors, except chromosome 3 illustrated in turquoise, unplaced scaffolds in tan, and scaffold 2 in misty rose. The points represent the ACs, and the labels are the genes residing in the DA; if a DAC entails more than one gene, then the genes are separated by a comma. The point size represents the length of DACs with a padj < 0.1, and PSI-BLAST found the gene’s symbol against *D. melanogaster*.

### Greater accessibility in hybrid identifying with *D. nasuta*

DESeq2 found 149 DAC regions, constituting more than 200Kb, of which 7 (∼32Kb) and 142 (∼169Kb) are highly accessible in hybrids identifying with *D. albomicans* and *D. nasuta,* respectively (**Fig. 1B and Supplemental Table S6)**. Similarly, edgeR identified 98 DAC regions spanning ∼130Kb (**Supplemental Table S7)**, where 93 (∼124 Kb) and 5 (∼22 Kb) regions are highly accessible in hybrids identifying with *D. nasuta* and *D. albomicans,* respectively (**Supplemental Fig. 6**). We have selected DESeq2 results for the subsequent analysis. Out of 149 DACs, 97 DACs (∼ 85Kb) occur in genomic regions which lack annotated genes. As a result, 52 DACs host the genes, albeit 48 genes, as the gene length varies, and in some cases, a single DAC hosts more than one gene (**Fig. 2**). The size of the DACs varies, and Muller F hosts a few of the largest DACs; the largest contiguous DAC is 41kb and contains three annotated genes: *Tdg*, *Pur-*α, and *CG32006*; where *Tdg* and *Pur-*α are involved in DNA mismatch repairing.

There are 24 DAC regions with more than a three-fold change, where 11 of them are associated with unplaced scaffolds, 10 of them with Muller F, and chr2L and chr3 have one each. But none of the 21 DACs in Muller F and unplaced scaffolds contain annotated genes; though chr2L does have one annotated gene, it is uncharacterized (see **Fig. 2B)**. A single DAC region of chr3 contains *Sugb* (Sugar baby) and has higher accessibility in hybrid identifying with *D. nasuta* (see **Fig. 2C**). Genes such as *Ten-a* (Tenascin accessory) and *beat-Ib* with roles in axon guidance also have higher accessibility in *D. nasuta* (see **Fig. 2A** and **2B**). The gene *qless* involved in the synthesis of the isoprenoid side chain of Coenzyme Q is also more accessible in hybrid identifying with *D. nasuta* (see **Fig. 2E**). Only one DAC region with > 1 fold-change has higher accessibility in hybrid identifying with *D. albomicans* and contains a long non-coding RNA (lncRNA) and the centrosomal protein-encoding gene *CG9170* (see **Fig. 2A**). The DACs also harbored nine long non-coding RNA (lncRNA); Muller F has four. These observations showed that the hybrid identifying with *D. nasuta* had a higher magnitude and a larger accessible genomic fraction, with 95.3% of all DAC found by DESeq2.

### DACs overlap the high F_ST_ region in Muller F

The overall autosomal genetic differentiation between populations of the same species (Mean F_ST_ ranges: 0.1324-0.3971) was much less than between-species differentiation (Mean F_ST_ ranges: 0.37-0.50) (**Fig. 3A, Supplemental Fig. 6, and Supplemental Fig. 7**). The within-species comparisons in *D*. *albomicans* have relatively higher levels of autosomal differentiation (Mean F_ST_ ranges: 0.15-0.42) than *D. nasuta* (Mean F_ST_ ranges: 0.18-0.22) (**Supplemental Fig. 7 and Supplemental Table S9**). Muller F showed the highest differentiation among the autosomes in between-species comparisons (Mean F_ST_ ranges: 0.5346-0.6162). However, Muller F retains the genome-wide pattern of lower differentiation between *D. nasuta* populations (Mean F_ST_ ranges: 0.1980-0.2542) compared to *D .albomicans* populations (Mean F_ST_ ranges: 0.1235-0.4041) (**Figure S7 and Supplemental Table S9**). Differentiation along Muller F is heterogeneous, with the initial 1.9 Mb having an F_ST_ < 0.3 and the subsequent region having an F_ST_ > 0.75.

**Figure 3:**
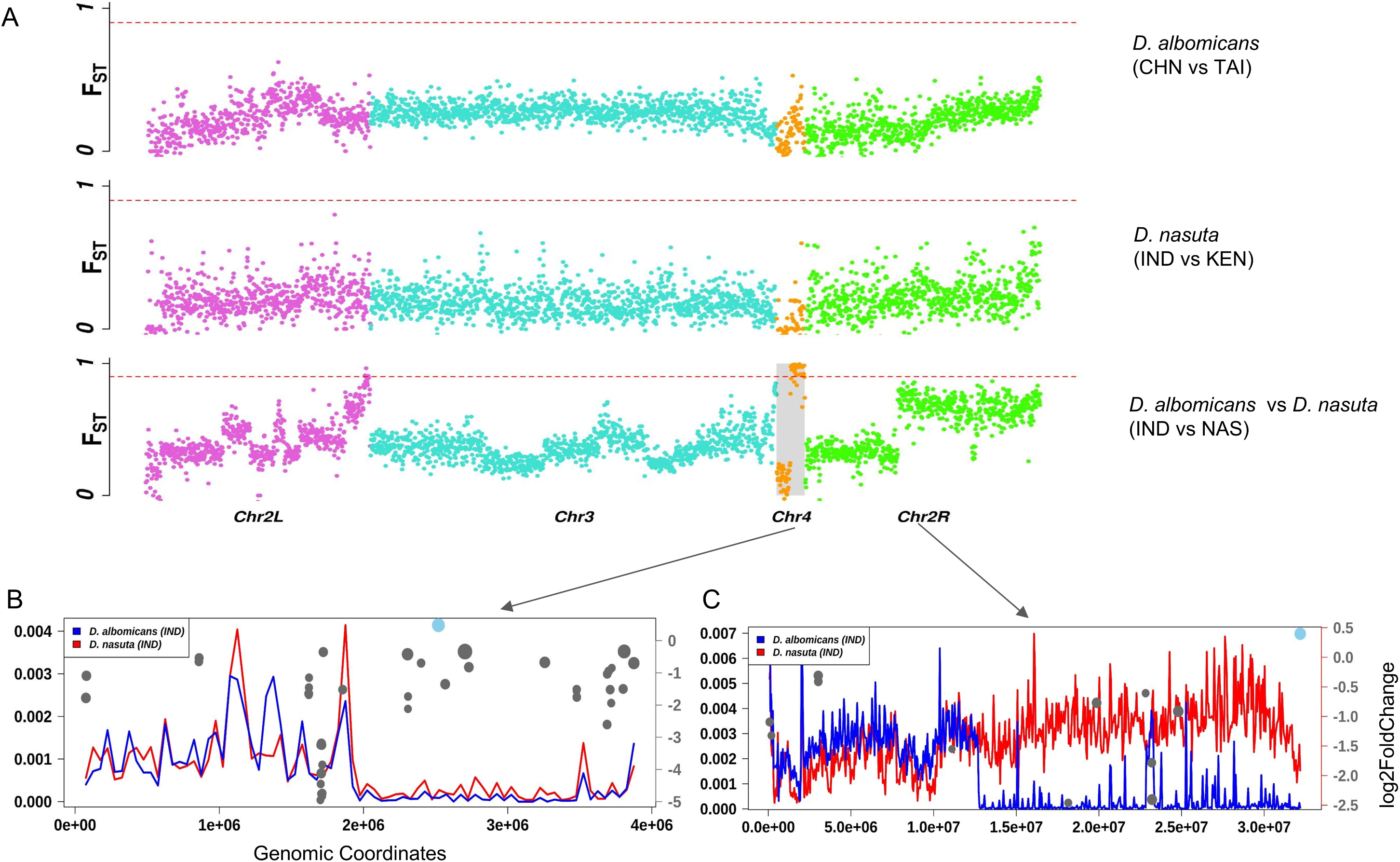
Genomic landscape of differentiation in *D. albomicans* and *D. nasuta* populations. **A** Pairwise genetic differentiation (FST) in 50-kb non-overlapping windows across the genome between *D. albomicans* population pair (CHN vs TAI) in top panel, between *D. nasuta* population pair (IND vs KEN) in second panel and between *D. albomicans* (IND) and *D. nasuta* (IND) population pair in third panel. Colours represent respective Muller elements; horizontal dotted red line demarcates FST estimate of 0.9. Chromosome 4 in third panel is highlighted in grey to show high differentiation region in the later half of chromosome. **B** Zoomed in view of the chromosome 4 for the same population pair of *D. albomicans* and *D. nasuta* to show their levels of genetic diversity depicted here as Watterson’s theta (0w) with blue and black lines respectively. The red dots denotes significant differential accessibility regions, whereas size of the dots is proportional to the length of the region. Most of the differential accessible regions overlap with high differentiation and low genomic diversity regions in later half of the chromosome. **C** Zoomed in view of the chromosome 2R for the same population pair of *D. albomicans* and *D. nasuta* to show their levels of genetic diversity depicted here as Watterson’s theta (0w) with blue and black lines respectively. The red and blue dots denotes significant differential accessibility regions, whereas size of the dots is proportional to the length of the region. Unlike chromosome 4, there is a drastic difference in genetic diversity levels between two species.

We identified several overlaps between the differential chromatin accessibility landscape and the high between-species genomic differentiation regions mostly occurring on Muller F (see **Supplementary Figure S6**). The highly differentiated distal end of Muller F has reduced genetic diversity (0w) in both *D*. *albomicans* and *D. nasuta* (see **Fig. 3B**). In contrast to Muller F, the differentiation peak on chr2R has a lower genetic diversity (0w) in hybrid identifying with *D*. *albomicans* compared to hybrid identifying with *D. nasuta* (see **Fig. 3C**). Numerous DAC regions, including the longest overlap with the island of elevated genomic differentiation at the end of Muller F. To ensure that these results are biologically meaningful, we excluded the repeat regions, and the results remained unchanged. Moreover, we found that the distal end of Muller F has a lower repeat density and higher exon density than the initial part (**Supplementary Figure S8**).

### Greater affinity towards pioneer TFs in hybrid identifying with *D. nasuta*

TOBIAS identified TFBS regions for 150 TF motif profiles across peak regions between two species. We found that TFs occupy 4.8% of the accessible genome across two species, where TFs occupy 3.22% in hybrid identifying with *D. albomicans*, 3.20% in hybrid identifying with *D. nasuta*, and 1.6% region in either one of the hybrids. TOBIAS calculated the TF activity score across two species and showed variation in the number of TF-occupied regions (**Table 2** and **Supplemental Table S10, S22**). The chrX had the fewest TF-bound sites than the Muller F (**Table 3** and **Supplemental Table S11**), explaining their lowest accessibility (**Figure 1B** and **Table 1**).

We found the highest TF activity scores for 46 TF (**Figure 4A)** and differential binding scores across hybrids (**Figure 4B**). Furthermore, hybrid identifying with *D. albomicans* showed higher binding affinity towards 28 TFs; hb, CG11294, CG32105, abd-A, al, H2.0, HGTX, lab, Lim1, otp, repo, unpg, zen2, mirr, br, C15, lms, Hmx, NK7.1, hbn, slou, CG11085, CG15696, CG32532, CG34031, unc-4, HHEX, and ubx. Whereas hybrid identifying with *D. nasuta* preferred eight TFs; Trl, Clamp, btd, odd, opa, Cf2, twi, and zld: a pioneer TF zld (Duan et al. 2021) chromatin remodelers; Trl and Clamp. The TF-TF network of 118 TFs (**Figure 4C and 4D**) shows that the higher affinity towards a specific TF changes the number of interacting partners. For example, the Trl network in hybrid identifying with *D. albomicans* and *D. nasuta* show variation in unique partners and the number of partners.

**Figure 4:**
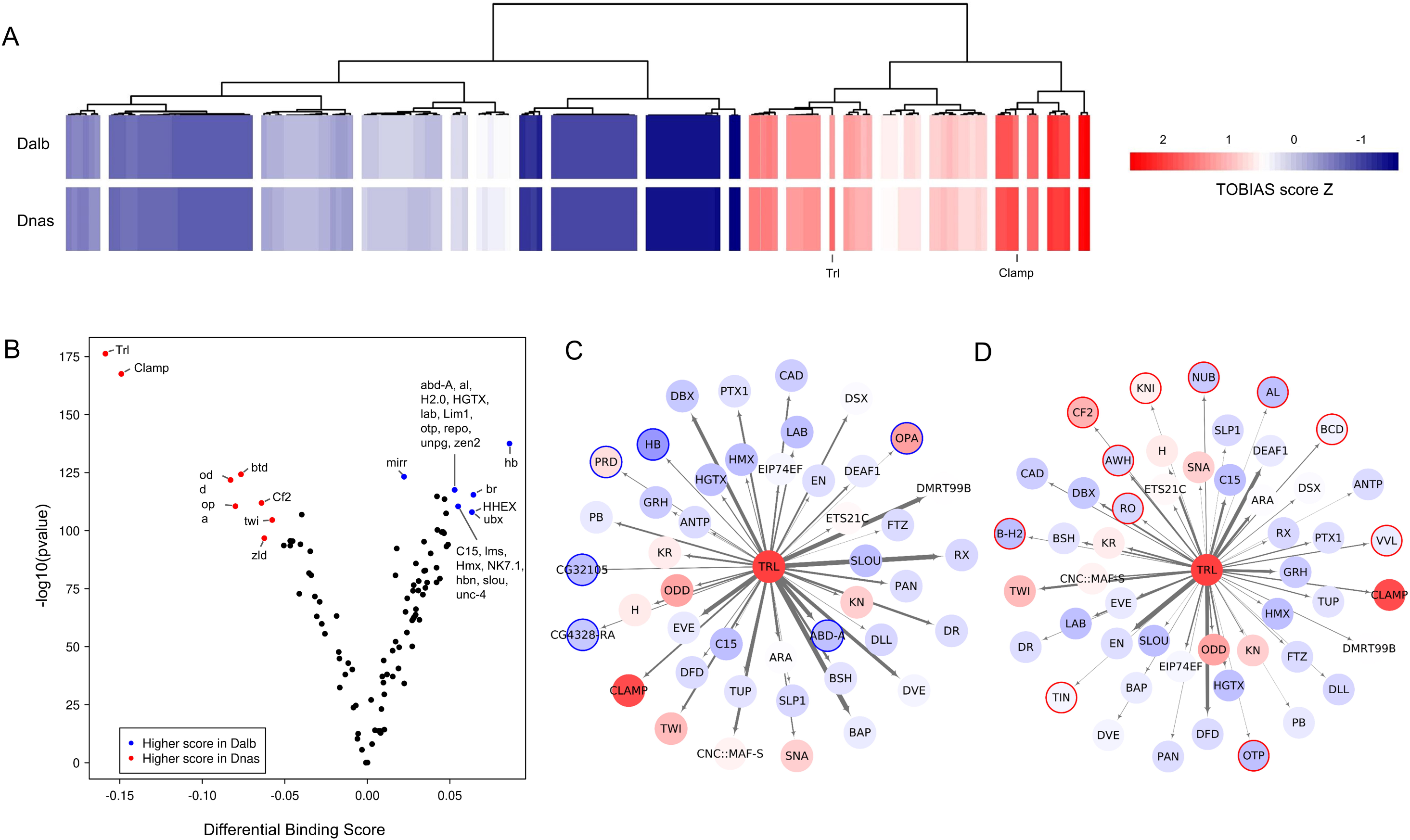
Transcription factor (TF) motifs across two species. **A** Heatmap of TF activity score across two hybrids, the scores were transformed into Z-score, where each row represents a TF in a species. Blue indicates low activity, and red indicates the higher TFs **B** TF motif activity score in testes across two species, by TOBIAS. Each dot in the volcano plot represents the activity scores of the TF motifs. The points were colored and labeled when the score meets the -log10(p-value) above 95% quantile and binding scores smaller/larger than the 5% and 95% quantiles **C** and **D** TF-TF network Trl in hybrids identifying with *D. albomicans* and *D. nasuta,* respectively. The color of node/TFs represents the binding scores of individual TFs, and the gradient of blue and red represents the higher binding score in hybrids identifying with *D. albomicans* and *D. nasuta,* respectively. The border colors around TFs show that they are unique to the individual species, and the edges/connection width represents the weights calculated by TOBIAS.

## Discussion

For the first time, we describe the former sex-chromosome – Muller F (Vicoso and Bachtrog 2013), driving the differential accessibility between hybrids of the *nasuta-albomicans* complex of *Drosophila*: *D. albomicans* and *D. nasuta.* There are few studies on the reversal of sex-chromosomes to autosomes, an unprecedented event in most organisms (Landeen and Presgraves 2013). Our study provides evidence for the existence of species that shows the transition of Muller F from sex-chromosome-like to autosome-like behavior in spermatogenesis. We found that hybrid identifying with *D. nasuta* has higher accessibility, though hybrid identifying with *D. albomicans* has lesser accessibility at the Muller F, consistent with another study (Vicoso and Bachtrog 2013). Our results showed a consistent pattern, where Muller F of hybrid identifying with *D. nasuta* is distinct from hybrid identifying with *D. albomicans* and other *Drosophila* species. For instance, TFBS prediction between hybrids implies hybrid identifying with *D. nasuta* binds differentially to the pioneer TFs, thus supporting the accessibility difference. Additionally, Muller F has higher genetic differentiation than other autosomes studied, hinting at the presence of genomic islands of differentiation.

### The nasuta-albomicans complex of Drosophila

Even though *D. albomicans* and *D. nasuta* are morphologically indistinguishable, few features set them apart. For instance, CRs (Ranganath and Hfigele 1982), followed by heterochromatin (Ranganath and Hfigele 1982; Wei et al. 2022), and later repeats and transposon density (Wei et al. 2022). A plethora of reasons recommends the consideration of testis in a variety of studies across species (Clark et al. 2007; Zhao et al. 2014; Xia et al. 2020; Witt et al. 2019). Furthermore, they are among the most studied adult tissue systems across species, especially in molecular and evolutionary biology. And are known to be highly transcriptionally active compared to any tissue of an organism (Witt et al. 2019). Our study shows greater accessibility across the genome, as expected from a highly dynamic tissue of an organism, more than 80% in all except one – least accessible chrX. Notably, the chrX undergoes inactivation during spermatogenesis, as illustrated by many studies (Hense et al. 2007; Richler et al. 1992; Turner 2007; Royo et al. 2010). Even though DACs are present across the genome, they nexus on a specific chromosome, Muller F (**Fig. 1B and C**). Even though we set out to study the CRs’ role in changing the chromatin accessibility landscape, we were intrigued by Muller F because of their compelling presence in DACs.

Muller F is one of the peculiar chromosomes found in *Drosophila.* They are primarily composed of heterochromatin, hosting less number of genes due to their petite size and lengthier genes; they enlist a vast number of repeats and a higher density of transposons (Funikov et al. 2020; Leung et al. 2015; Wei et al. 2022). For example, distal regions of Muller F of *D. albomicans,* 1.9Mb, showed a higher gene density, 113 genes with an average length of 10.8Kb. Additionally, Muller F’s idiosyncratic lineage of evolution (Vicoso and Bachtrog 2013) piques interest in exploring why they host nearly 50% of all identified DACs (**Fig. 1B and C**) and the functional role of the genes it contains. Muller F, in our study, hosts one of the largest DAC with a three-fold difference between hybrids identifying with *D. albomicans* and *D. nasuta*, entailing 41kb of 3.9Mb of its length, and has three annotated genes: *Tdg*, *Pur-*α, and *CG32006*. Recent studies have observed that the *D. nasuta* accommodate high repeat contents compared to *D. albomicans* (Wei et al. 2022), and it’s known that the repeats elicit the DNA mismatches (Li et al. 2002; Iyer et al. 2015), repaired by the enzymes like *Tdg* and *Pur-*α (Hardeland et al. 2003; White et al. 2009). This information led us to speculate that the repeats load and the repairing genes could explain the Muller F three-fold differential accessibility.

This observation (**Fig. 1B and C**) led us to suspect that the Muller F could harbor species’ genomic islands of differentiation and potentially be a reproductive barrier (Feder et al. 2012). To discern our speculation, we scanned for genome-wide divergence in public datasets of the species (**Supplemental Table S2**). The F_ST_ across the chromosomes showed few regions across the genome, chr2L, chr3, Muller F, chr2R, and chrX; generally, sex chromosomes exhibit high divergence (Wolf and Ellegren 2017). All these regions have higher F_ST_; though sex chromosomes show high divergence, only the distal end of the Muller F showed fixed divergence (**Fig. 3A, B, and C**), and they also juxtaposed with the DACs in our study. We have detected differentiation across various genome regions, hinting that these species might be experiencing low or no gene flow, a characteristic of allopatric species (Feder et al. 2012).

After assessing the DAC and fixed divergence across two hybrids, we want to identify if these elements affect the TF binding. Indeed, we found differences, specifically in hybrid identifying with *D. nasuta*, involving two pioneer TFs: ZLD and CLAMP. Pioneer TFs are the only TFs that can bind to nucleosomal DNA, and modulate for chromatin accessibility, thus associated with open chromatin and active transcription (McPherson et al. 1993; Zaret 2020). ZLD and CLAMP are the drivers of Drosophila zygotic genome activation (ZGA), and ZLD is necessary for hundreds of regions’ open chromatin (Schulz et al. 2015). Thus, pioneer TFs contribute towards higher accessibility observed in hybrid identifying with *D. nasuta*. Additionally, hybrid identifying with *D. nasuta* also has a higher binding affinity for TRL or GAF (Trithorax-like), which belongs to Trithorax (TrxG) group proteins. TrxG and Polycomb group (PcG) can regulate their target at multiple levels, modifying the local chromatin to higher chromatin order and global genome structure (Schuettengruber et al. 2017). TRL is involved in modifying the accessibility of promoters, allowing other TFs to bind and start transcription (Farkas et al. 1994). We believe that the accessibility changes seen in hybrid identifying with *D. nasuta* are a culmination of the higher activity of these TFs.

### Limitations

During the analysis of the ATAC-seq data using different reference genomes, it was noticed that differential mapping bias towards the parental species affects specific genomic regions. Since the hybrids used in this study have differing levels of hybrid ancestry (see **Supplementary Text**), disentangling the effect of mapping bias from biological differences in accessibility remains challenging. Future studies with high-quality reference genomes and sophisticated tools to overcome the challenges of mapping bias in hybrids will be crucial for further progress. Although the mapping bias observed could be responsible for some of the biological patterns observed, we can rule out the existence of a distinctive biological signal. Therefore, we urge caution in the interpretation of these results.

## Conclusion

We set out to understand the interplay between epigenetics, chromosomal rearrangement, and speciation using ATAC-seq and DNA-seq of members of the *nasuta-albomicans* complex of *Drosophila* and hybrids. Our results showed chromatin accessibility differentiation, one of the many components of epigenetics, between the two hybrids. Though we found few DACs in the Robertsonian fused chromosome, Muller F, one of the smallest assembled chromosomes, drew our attention. Due to the density of DACs identified and because it’s one of the notable karyotypic differences found in *D. albomicans* and *D. nasuta*. *D. albomicans* hosts a lengthier and heterochromatinized version of Muller F than one in *D. nasuta*. The result reemphasizes the observation with the juxtaposed region of significantly higher accessibility and genetic differentiation in hybrid identifying with *D. nasuta.* The higher accessibility in hybrid identifying with *D. nasuta* is probably due to their strong affinity towards binding pioneer TFs and chromatin modifiers, like CLAMP, TRL, and ZLD. Our main objective was to understand how CRs and ACs affect the speciation in general and precisely how they shape the genome of hybrids of closely related species, *D. albomicans* and *D. nasuta*. Our findings conclude that the first difference that could arise in heterochromatin. We urge future studies to carefully observe the interaction of karyotype, diversity of accessibility and genetic variation, and TF motifs to understand the speciation.

## Supporting information

Review from journal 1

Review from journal 2

## Note to Journals

If any peer-reviewed journal is interested in this manuscript, they are requested to email the authors. This manuscript has already been reviewed by two journals and rejected without an option to resubmit. The detailed reviewer comments (without journal names) are provided as Supplementary Files. We hope these reviews provide the required background information.

## Authors contributions

R.P. designed the study, obtained funding and performed all the experiments. R.P. wrote the manuscript with inputs from A.B.P and N.V. R.P. analyzed all the ATAC-seq data. A.B.P performed the population genetic analysis and provided crucial inputs on the integration of diverse analysis. All authors reviewed the manuscript.

## Data accessibility

The associated data is available in an easy-to-view format on the GitHub repository: https://github.com/RadPa/Chromatin-accessibility-of-hybrids-of-D.-albomicans-and-D.-nasuta.git.

## References

Abzhanov A, Protas M, Grant BR, Grant PR, Tabin CJ. 2004. Bmp4 and Morphological Variation of Beaks in Darwin’s Finches. Science (80-) 305: 1462–1465. www.science.org/cgi/content/full/305/5689/1457/ (Accessed March 10, 2022).

Allis CD, Jenuwein T. 2016. The molecular hallmarks of epigenetic control. Nat Rev Genet 2016 178 17: 487–500. https://www.nature.com/articles/nrg.2016.59 (Accessed May 3, 2022).

Allshire RC, Madhani HD. 2018. Ten principles of heterochromatin formation and function. Nat Rev Mol Cell Biol 19: 229–244. https://www.nature.com/articles/nrm.2017.119 (Accessed March 12, 2022).

Amemiya HM, Kundaje A, Boyle AP. 2019. The ENCODE Blacklist: Identification of Problematic Regions of the Genome. Sci Rep 9: 9354. https://www.nature.com/articles/s41598-019-45839-z (Accessed October 27, 2021).

Andrews S, others. 2010. FastQC: a quality control tool for high throughput sequence data. 2010. Https://WwwBioinformaticsBabrahamAcUk/Projects/Fastqc/ http://www.bioinformatics.babraham.ac.uk/projects/. https://www.bioinformatics.babraham.ac.uk/projects/fastqc/ (Accessed November 20, 2021).

Ayala FJ, Coluzzi M. 2005. Chromosome speciation: Humans, Drosophila, and mosquitoes. Proc Natl Acad Sci 102: 6535–6542. http://www.pnas.org/cgi/doi/10.1073/pnas.0501847102 (Accessed August 24, 2019).

Balsalobre A, Drouin J. 2022. Pioneer factors as master regulators of the epigenome and cell fate. Nat Rev Mol Cell Biol 2022 1–16. https://www.nature.com/articles/s41580-022-00464-z (Accessed May 22, 2022).

Bentsen M, Goymann P, Schultheis H, Klee K, Petrova A, Wiegandt R, Fust A, Preussner J, Kuenne C, Braun T, et al. 2020. ATAC-seq footprinting unravels kinetics of transcription factor binding during zygotic genome activation. Nat Commun 11: 4267. https://www.nature.com/articles/s41467-020-18035-1 (Accessed August 18, 2021).

Bhutkar A, Schaeffer SW, Russo SM, Xu M, Smith TF, Gelbart WM. 2008. Chromosomal Rearrangement Inferred From Comparisons of 12 Drosophila Genomes. Genetics 179: 1657–1680. https://academic.oup.com/genetics/article/179/3/1657/6063947 (Accessed October 25, 2019).

Burri R, Nater A, Kawakami T, Mugal CF, Olason PI, Smeds L, Suh A, Dutoit L, Bureš S, Garamszegi LZ, et al. 2015. Linked selection and recombination rate variation drive the evolution of the genomic landscape of differentiation across the speciation continuum of Ficedula flycatchers. Genome Res 25: 1656–1665. https://genome.cshlp.org/content/25/11/1656.full (Accessed April 28, 2022).

Castro-Mondragon JA, Riudavets-Puig R, Rauluseviciute I, Lemma RB, Turchi L, Blanc-Mathieu R, Lucas J, Boddie P, Khan A, As Manosalva P, Erez N, et al. 2022. JASPAR 2022: the 9th release of the open-access database of transcription factor binding profiles. Nucleic Acids Res 50: D165–D173. https://academic.oup.com/nar/article/50/D1/D165/6446529 (Accessed February 17, 2022).

Chan YF, Marks ME, Jones FC, Villarreal G, Shapiro MD, Brady SD, Southwick AM, Absher DM, Grimwood J, Schmutz J, et al. 2010. Adaptive Evolution of Pelvic Reduction in Sticklebacks by Recurrent Deletion of a Pitx1 Enhancer. Science (80-) 327: 302–305. https://www.science.org/doi/10.1126/science.1182213 (Accessed March 10, 2022).

Clark AG, Eisen MB, Smith DR, Bergman CM, Oliver B, Markow TA, Kaufman TC, Kellis M, Gelbart W, Iyer VN, et al. 2007. Evolution of genes and genomes on the Drosophila phylogeny. Nature 450: 203–218. https://www.nature.com/articles/nature06341 (Accessed February 17, 2022).

Comeron JM, Ratnappan R, Bailin S. 2012. The Many Landscapes of Recombination in Drosophila melanogaster ed. D.A. Petrov. PLoS Genet 8: e1002905. https://journals.plos.org/plosgenetics/article?id=10.1371/journal.pgen.1002905 (Accessed April 28, 2022).

Corbett-Detig RB, Hartl DL. 2012. Population Genomics of Inversion Polymorphisms in Drosophila melanogaster. PLOS Genet 8: e1003056. https://journals.plos.org/plosgenetics/article?id=10.1371/journal.pgen.1003056 (Accessed March 12, 2022).

Dainat J, Hereñú D, pascal-git. 2021. NBISweden/AGAT: AGAT-v0.8.0. https://zenodo.org/record/5336786 (Accessed October 27, 2021).

de Vos JM, Augustijnen H, Bätscher L, Lucek K. 2020. Speciation through chromosomal fusion and fission in Lepidoptera. Philos Trans R Soc B Biol Sci 375: 20190539. https://royalsocietypublishing.org/doi/abs/10.1098/rstb.2019.0539 (Accessed September 28, 2021).

Duan JE, Rieder LE, Colonnetta MM, Huang A, McKenney M, Watters S, Deshpande G, Jordan WT, Fawzi NL, Larschan EN. 2021. Clamp and zelda function together to promote drosophila zygotic genome activation. Elife 10.

Elgin SCR, Reuter G. 2013. Position-Effect Variegation, Heterochromatin Formation, and Gene Silencing in Drosophila. Cold Spring Harb Perspect Biol 5: a017780–a017780. http://cshperspectives.cshlp.org/lookup/doi/10.1101/cshperspect.a017780 (Accessed October 25, 2019).

Ewels P, Magnusson M, Lundin S, Käller M. 2016. MultiQC: summarize analysis results for multiple tools and samples in a single report. Bioinformatics 32: 3047–3048. https://academic.oup.com/bioinformatics/article/32/19/3047/2196507 (Accessed October 27, 2021).

Farkas G, Gausz J, Galloni M, Reuter G, Gyurkovics H, Karch F. 1994. The Trithorax-like gene encodes the Drosophila GAGA factor. Nature 371: 806–808. https://pubmed.ncbi.nlm.nih.gov/7935842/ (Accessed February 23, 2022).

Feder JL, Egan SP, Nosil P. 2012. The genomics of speciation-with-gene-flow. Elsevier Current Trends https://www.sciencedirect.com/science/article/pii/S0168952512000492 (Accessed October 1, 2019).

Funikov SY, Rezvykh AP, Kulikova DA, Zelentsova ES, Protsenko LA, Chuvakova LN, Tyukmaeva VI, Arkhipova IR, Evgen’ev MB. 2020. Adaptation of gene loci to heterochromatin in the course of Drosophila evolution is associated with insulator proteins. Sci Rep 10: 11893. https://www.nature.com/articles/s41598-020-68879-2 (Accessed September 1, 2021).

Gibilisco L, Zhou Q, Mahajan S, Bachtrog D. 2016. Alternative Splicing within and between Drosophila Species, Sexes, Tissues, and Developmental Stages ed. G.S. Barsh. PLOS Genet 12: e1006464. https://journals.plos.org/plosgenetics/article?id=10.1371/journal.pgen.1006464 (Accessed November 17, 2021).

Goettel W, Messing J. 2010. Divergence of gene regulation through chromosomal rearrangements. BMC Genomics 11: 1–19. https://bmcgenomics.biomedcentral.com/articles/10.1186/1471-2164-11-678 (Accessed November 20, 2021).

Guerrero RF, Kirkpatrick M. 2014. LOCAL ADAPTATION AND THE EVOLUTION OF CHROMOSOME FUSIONS. Evolution (N Y) 68: 2747–2756. https://onlinelibrary.wiley.com/doi/full/10.1111/evo.12481 (Accessed November 8, 2021).

Hardeland U, Bentele M, Jiricny J, Schär P. 2003. The versatile thymine DNA-glycosylase: A comparative characterization of the human, Drosophila and fission yeast orthologs. Nucleic Acids Res 31: 2261–2271. https://academic.oup.com/nar/article/31/9/2261/1080396 (Accessed May 27, 2022).

Harewood L, Fraser P. 2014. The impact of chromosomal rearrangements on regulation of gene expression. Hum Mol Genet 23: R76–R82. https://academic.oup.com/hmg/article/23/R1/R76/2900848 (Accessed September 28, 2021).

Hense W, Baines JF, Parsch J. 2007. X Chromosome Inactivation during Drosophila Spermatogenesis ed. M.A.F. Noor. PLoS Biol 5: e273. https://journals.plos.org/plosbiology/article?id=10.1371/journal.pbio.0050273 (Accessed February 7, 2022).

Hollox EJ, Zuccherato LW, Tucci S. 2022. Genome structural variation in human evolution. Elsevier http://www.cell.com/article/S0168952521001852/fulltext (Accessed March 12, 2022).

Holsinger KE, Weir BS. 2009. Genetics in geographically structured populations: defining, estimating and interpreting FST. Nat Rev Genet 10: 639–650. https://www.nature.com/articles/nrg2611 (Accessed March 12, 2022).

Holtzman L, Gersbach CA. 2018. Editing the epigenome: Reshaping the genomic landscape. Annu Rev Genomics Hum Genet 19: 43–71. https://www.annualreviews.org/doi/abs/10.1146/annurev-genom-083117-021632 (Accessed May 3, 2022).

Iyer RR, Pluciennik A, Napierala M, Wells RD. 2015. DNA Triplet Repeat Expansion and Mismatch Repair. Annu Rev Biochem 84: 199–226. https://www.annualreviews.org/doi/abs/10.1146/annurev-biochem-060614-034010 (Accessed May 31, 2022).

Janssen A, Colmenares SU, Karpen GH. 2018. Heterochromatin: Guardian of the Genome. Annu Rev Cell Dev Biol 34: 265–288. 10.1146/annurev-cellbio-100617- (Accessed March 11, 2022).

Karimzadeh M, Ernst C, Kundaje A, Hoffman MM. 2018. Umap and Bismap: quantifying genome and methylome mappability. Nucleic Acids Res 46: e120–e120. https://academic.oup.com/nar/article/46/20/e120/5086676 (Accessed October 27, 2021).

Kirkpatrick M, Barton N. 2006. Chromosome Inversions, Local Adaptation and Speciation. Genetics 173: 419–434. https://academic.oup.com/genetics/article/173/1/419/6061572 (Accessed March 12, 2022).

Kitagawa O, Wakahama K-I, Fuyama Y, Shimada Y, Takanashi E, Hatsumi M, Uwabo M, Mita Y. 1982. Genetic studies of the Drosophila nasuta subgroup, with notes on distribution and morphology. Japanese J Genet 57: 113–141. http://www.jstage.jst.go.jp/article/ggs1921/57/2/57_2_113/_article (Accessed November 16, 2021).

Klemm SL, Shipony Z, Greenleaf WJ. 2019. Chromatin accessibility and the regulatory epigenome. Nature Publishing Group www.nature.com/nrg (Accessed October 17, 2019).

Kondili M, Fust A, Preussner J, Kuenne C, Braun T, Looso M. 2017. UROPA: A tool for Universal RObust Peak Annotation. Sci Rep 7: 1–12. https://www.nature.com/articles/s41598-017-02464-y (Accessed February 17, 2022).

Korneliussen TS, Albrechtsen A, Nielsen R. 2014. ANGSD: Analysis of Next Generation Sequencing Data. BMC Bioinformatics 15. https://bmcbioinformatics.biomedcentral.com/articles/10.1186/s12859-014-0356-4 (Accessed October 27, 2021).

Kou Y, Liao Y, Toivainen T, Lv Y, Tian X, Emerson JJ, Gaut BS, Zhou Y. 2020. Evolutionary Genomics of Structural Variation in Asian Rice (Oryza sativa) Domestication. Mol Biol Evol 37: 3507–3524. https://academic.oup.com/mbe/article/37/12/3507/5873527 (Accessed March 12, 2022).

Landeen EL, Presgraves DC. 2013. Evolution: From autosomes to sex chromosomes - And back. Curr Biol 23: R848–R850. https://linkinghub.elsevier.com/retrieve/pii/S0960982213010270 (Accessed June 2, 2022).

Langmead B, Trapnell C, Pop M, Salzberg SL. 2009. Ultrafast and memory-efficient alignment of short DNA sequences to the human genome. Genome Biol 10: R25. https://genomebiology.biomedcentral.com/articles/10.1186/gb-2009-10-3-r25 (Accessed October 27, 2021).

Lee CK, Shibata Y, Rao B, Strahl BD, Lieb JD. 2004. Evidence for nucleosome depletion at active regulatory regions genome-wide. Nat Genet 36: 900–905. https://www.nature.com/articles/ng1400 (Accessed March 11, 2022).

Leung W, Shaffer CD, Reed LK, Smith ST, Barshop W, Dirkes W, Dothager M, Lee P, Wong J, Xiong D, et al. 2015. Drosophila Muller F Elements Maintain a Distinct Set of Genomic Properties Over 40 Million Years of Evolution. G3 Genes|Genomes|Genetics 5: 719–740. http://www.g3journal.org/lookup/suppl/doi:10.1534/g3.114.015966/-/DC1 (Accessed November 25, 2020).

Li H, Durbin R. 2009. Fast and accurate short read alignment with Burrows–Wheeler transform. Bioinformatics 25: 1754–1760. https://academic.oup.com/bioinformatics/article/25/14/1754/225615 (Accessed November 20, 2021).

Li H, Handsaker B, Wysoker A, Fennell T, Ruan J, Homer N, Marth G, Abecasis G, Durbin R. 2009. The Sequence Alignment/Map format and SAMtools. Bioinformatics 25: 2078–2079. https://academic.oup.com/bioinformatics/article/25/16/2078/204688 (Accessed October 27, 2021).

Li YC, Korol AB, Fahima T, Beiles A, Nevo E. 2002. Microsatellites: Genomic distribution, putative functions and mutational mechanisms: A review. Mol Ecol 11: 2453–2465. https://onlinelibrary.wiley.com/doi/full/10.1046/j.1365-294X.2002.01643.x (Accessed May 31, 2022).

Love MI, Huber W, Anders S. 2014. Moderated estimation of fold change and dispersion for RNA-seq data with DESeq2. Genome Biol 15: 550. https://genomebiology.biomedcentral.com/articles/10.1186/s13059-014-0550-8 (Accessed October 27, 2021).

Lukhtanov VA, Dincă V, Talavera G, Vila R. 2011. Unprecedented within-species chromosome number cline in the Wood White butterfly Leptidea sinapis and its significance for karyotype evolution and speciation. BMC Evol Biol 11: 109. https://bmcecolevol.biomedcentral.com/articles/10.1186/1471-2148-11-109 (Accessed November 16, 2021).

Mai D, Bachtrog D. 2021. Molecular characterization of inversion breakpoints in the Drosophila nasuta species group. bioRxiv 2021.06.01.446624. https://www.biorxiv.org/content/10.1101/2021.06.01.446624v1 (Accessed March 13, 2022).

Martin M. 2011. Cutadapt removes adapter sequences from high-throughput sequencing reads. EMBnet.journal 17: 10. https://github.com/marcelm/cutadapt (Accessed November 20, 2021).

Mathers TC, Wouters RHM, Mugford ST, Swarbreck D, van Oosterhout C, Hogenhout SA. 2021. Chromosome-Scale Genome Assemblies of Aphids Reveal Extensively Rearranged Autosomes and Long-Term Conservation of the X Chromosome ed. A. Ouangraoua. Mol Biol Evol 38: 856–875. https://academic.oup.com/mbe/article/38/3/856/5910553 (Accessed November 16, 2021).

McClintock B. 1953. Induction of Instability at Selected Loci in Maize. Genetics 38: 579–599. https://pubmed.ncbi.nlm.nih.gov/17247459/ (Accessed November 20, 2021).

McPherson CE, Shim EY, Friedman DS, Zaret KS. 1993. An active tissue-specific enhancer and bound transcription factors existing in a precisely positioned nucleosomal array. Cell 75: 387–398.

Muller HJ. 1930. Types of visible variations induced by X-rays inDrosophila. J Genet 22: 299–334. https://link.springer.com/article/10.1007/BF02984195 (Accessed November 20, 2021).

Navarro A, Barton NH. 2003. Chromosomal Speciation and Molecular Divergence--Accelerated Evolution in Rearranged Chromosomes. Science (80-) 300: 321–324. https://www.science.org/doi/abs/10.1126/science.1080600 (Accessed April 28, 2022).

Nozawa M, Minakuchi Y, Satomura K, Kondo S, Toyoda A, Tamura K. 2021. Shared evolutionary trajectories of three independent neo-sex chromosomes in Drosophila. Genome Res 31: 2069–2079. https://genome.cshlp.org/content/31/11/2069.full (Accessed November 17, 2021).

Ou J, Liu H, Yu J, Kelliher MA, Castilla LH, Lawson ND, Zhu LJ. 2018. ATACseqQC: a Bioconductor package for post-alignment quality assessment of ATAC-seq data. BMC Genomics 19: 169. https://bmcgenomics.biomedcentral.com/articles/10.1186/s12864-018-4559-3 (Accessed June 28, 2019).

Poelstra JW, Vijay N, Bossu CM, Lantz H, Ryll B, Müller I, Baglione V, Unneberg P, Wikelski M, Grabherr MG, et al. 2014. The genomic landscape underlying phenotypic integrity in the face of gene flow in crows. Science (80-) 344: 1410–1414.

Prud’homme B, Gompel N, Carroll SB. 2007. Emerging principles of regulatory evolution. Proc Natl Acad Sci 104: 8605–8612. www.nasonline.org/adaptation_and_complex_design. (Accessed March 10, 2022).

Quinlan AR, Hall IM. 2010. BEDTools: A flexible suite of utilities for comparing genomic features. Bioinformatics 26: 841–842. https://academic.oup.com/bioinformatics/article/26/6/841/244688 (Accessed May 30, 2019).

Ranganath HA, Hfigele K. 1982. The Chromosomes of Two Drosophila Races: D. nasuta nasuta and D. n. albomicana I. Distribution and Differentiation of Heterochromatin *. Chromosom 85: 83–92.

Richards EJ, Elgin SCR. 2002. Epigenetic Codes for Heterochromatin Formation and Silencing: Rounding up the Usual Suspects. Cell 108: 489–500.

Richler C, Soreq H, Wahrman J. 1992. X inactivation in mammalian testis is correlated with inactive X–specific transcription. Nat Genet 2: 192–195. https://www.nature.com/articles/ng1192-192 (Accessed February 17, 2022).

Rieseberg LH. 2001. Chromosomal rearrangements and speciation. Trends Ecol Evol 16: 351–358. http://www.cell.com/article/S0169534701021875/fulltext (Accessed September 27, 2021).

Robinson MD, McCarthy DJ, Smyth GK. 2010. edgeR: a Bioconductor package for differential expression analysis of digital gene expression data. Bioinformatics 26: 139–140. https://academic.oup.com/bioinformatics/article/26/1/139/182458 (Accessed February 17, 2022).

Royo H, Polikiewicz G, Mahadevaiah SK, Prosser H, Mitchell M, Bradley A, de Rooij DG, Burgoyne PS, Turner JMA. 2010. Evidence that Meiotic Sex Chromosome Inactivation Is Essential for Male Fertility. Curr Biol 20: 2117–2123. https://linkinghub.elsevier.com/retrieve/pii/S0960982210014351 (Accessed February 17, 2022).

Satomura K, Tamura K. 2016. Ancient Male Recombination Shaped Genetic Diversity of Neo-Y Chromosome in Drosophila albomicans. Mol Biol Evol 33: 367–374. https://academic.oup.com/mbe/article/33/2/367/2579414 (Accessed November 17, 2021).

Schuettengruber B, Bourbon H-M, Di Croce L, Cavalli G. 2017. Genome Regulation by Polycomb and Trithorax: 70 Years and Counting. Cell 171: 34–57. https://linkinghub.elsevier.com/retrieve/pii/S0092867417308905 (Accessed February 23, 2022).

Schulz KN, Bondra ER, Moshe A, Villalta JE, Lieb JD, Kaplan T, McKay DJ, Harrison MM. 2015. Zelda is differentially required for chromatin accessibility, transcription factor binding, and gene expression in the early Drosophila embryo. Genome Res 25: 1715–1726. https://genome.cshlp.org/content/25/11/1715.full (Accessed March 22, 2022).

Shannon P, Markiel A, Ozier O, Baliga NS, Wang JT, Ramage D, Amin N, Schwikowski B, Ideker T. 2003. Cytoscape: A Software Environment for Integrated Models of Biomolecular Interaction Networks. Genome Res 13: 2498–2504. https://genome.cshlp.org/content/13/11/2498.full (Accessed February 17, 2022).

Stewart NB, Rogers RL. 2019. Chromosomal rearrangements as a source of new gene formation in Drosophila yakuba ed. H.S. Malik. PLOS Genet 15: e1008314. https://journals.plos.org/plosgenetics/article?id=10.1371/journal.pgen.1008314 (Accessed September 1, 2021).

Sun F-L, Cuaycong MH, Elgin SCR. 2001. Long-Range Nucleosome Ordering Is Associated with Gene Silencing in Drosophila melanogaster Pericentric Heterochromatin. Mol Cell Biol 21: 2867–2879. /pmc/articles/PMC86916/ (Accessed March 11, 2022).

Tarbell ED, Liu T. 2019. HMMRATAC: a Hidden Markov ModeleR for ATAC-seq. Nucleic Acids Res 47: e91–e91. https://academic.oup.com/nar/article/47/16/e91/5519166 (Accessed October 27, 2021).

Turner JMA. 2007. Meiotic sex chromosome inactivation. Development 134: 1823–1831. https://journals.biologists.com/dev/article/134/10/1823/52756/Meiotic-sex-chromosome-inactivation (Accessed February 17, 2022).

Turner TL, Hahn MW, Nuzhdin S V. 2005. Genomic Islands of Speciation in Anopheles gambiae ed. N. Barton. PLoS Biol 3: e285. https://journals.plos.org/plosbiology/article?id=10.1371/journal.pbio.0030285 (Accessed April 28, 2022).

Van der Auwera GA, O’Connor BD. 2020. Genomics in the Cloud. https://www.oreilly.com/library/view/genomics-in-the/9781491975183/ (Accessed October 27, 2021).

Vicoso B, Bachtrog D. 2013. Reversal of an ancient sex chromosome to an autosome in Drosophila. Nature 499: 332–335. https://www.nature.com/articles/nature12235 (Accessed March 19, 2021).

Wei KH-C, Mai D, Chatla K, Bachtrog D. 2021. Dynamics and impacts of transposable element proliferation during the Drosophila nasuta species group radiation. bioRxiv 2021.08.12.456169. https://www.biorxiv.org/content/10.1101/2021.08.12.456169v1 (Accessed February 18, 2022).

Wei KH-C, Mai D, Chatla K, Bachtrog D. 2022. Dynamics and Impacts of Transposable Element Proliferation in the Drosophila nasuta Species Group Radiation ed. R. Rogers. Mol Biol Evol 39. https://academic.oup.com/mbe/article/39/5/msac080/6575839 (Accessed May 30, 2022).

Wei KHC, Bachtrog D. 2019. Ancestral male recombination in Drosophila albomicans produced geographically restricted neo-Y chromosome haplotypes varying in age and onset of decay ed. K.A. Dyer. PLOS Genet 15: e1008502. 10.1371/journal.pgen.1008502 (Accessed November 26, 2020).

White MK, Johnson EM, Khalili K. 2009. Multiple roles for Purα in cellular and viral regulation. Cell Cycle 8: 414–420. https://www.tandfonline.com/doi/abs/10.4161/cc.8.3.7585 (Accessed May 30, 2022).

Wilson FD, Wheeler MR, Harget M, Kambysellis M. 1969. Cytogenetic relations in the Drosophila nasuta subgroup of the immigrans group of species. Univ Texas Publ 6918: 207–253. https://ci.nii.ac.jp/naid/10015117093 (Accessed November 16, 2021).

Witt E, Benjamin S, Svetec N, Zhao L. 2019. Testis single-cell RNA-seq reveals the dynamics of de novo gene transcription and germline mutational bias in Drosophila. Elife 8. https://elifesciences.org/articles/47138 (Accessed August 2, 2021).

Wittkopp PJ, Haerum BK, Clark AG. 2008. Regulatory changes underlying expression differences within and between Drosophila species. Nat Genet 40: 346–350. https://www.nature.com/articles/ng.77 (Accessed March 10, 2022).

Wolf JBW, Ellegren H. 2017. Making sense of genomic islands of differentiation in light of speciation. Nat Rev Genet 18: 87–100. http://www.nature.com/articles/nrg.2016.133 (Accessed May 23, 2019).

Wray GA. 2003. The Evolution of Transcriptional Regulation in Eukaryotes. Mol Biol Evol 20: 1377–1419. https://academic.oup.com/mbe/article-lookup/doi/10.1093/molbev/msg140 (Accessed December 19, 2019).

Xia B, Yan Y, Baron M, Wagner F, Barkley D, Chiodin M, Kim SY, Keefe DL, Alukal JP, Boeke JD, et al. 2020. Widespread Transcriptional Scanning in the Testis Modulates Gene Evolution Rates. Cell 180: 248–262.e21. https://linkinghub.elsevier.com/retrieve/pii/S0092867419313777 (Accessed February 17, 2022).

Zaret KS. 2020. Pioneer Transcription Factors Initiating Gene Network Changes. Annu Rev Genet 54. /pmc/articles/PMC7900943/ (Accessed February 15, 2022).

Zhao L, Saelao P, Jones CD, Begun DJ. 2014. Origin and spread of de novo genes in Drosophila melanogaster populations. Science (80-) 343: 769–772. https://www.science.org (Accessed February 17, 2022).

Zhou Q, Zhu H, Huang Q, Zhao L, Zhang G, Roy SW, Vicoso B, Xuan Z, Ruan J, Zhang Y, et al. 2012. Deciphering neo-sex and B chromosome evolution by the draft genome of Drosophila albomicans. BMC Genomics 13: 109. https://link.springer.com/articles/10.1186/1471-2164-13-109 (Accessed November 17, 2021).

Zhu T, Liang C, Meng Z, Guo S, Zhang R. 2017. GFF3sort: a novel tool to sort GFF3 files for tabix indexing. BMC Bioinformatics 18: 482. https://bmcbioinformatics.biomedcentral.com/articles/10.1186/s12859-017-1930-3 (Accessed October 27, 2021).

