## Supplementary material for "Chromatin accessibility differences between the hybrids of *nasuta-albomicans* complex of *Drosophila*": Review from journal 1

Editor (Comments to the Author):

Thank you for submitting your manuscript to the journal. We appreciate the effort and thought that went into your work, which explores chromatin accessibility in hybrids of Drosophila albomicans and D. nasuta, and its potential role in reproductive isolation.

Both reviewers acknowledged the relevance and originality of the topic, particularly given the limited data available on epigenomic divergence in hybrid systems. However, after careful consideration, we regret to inform you that we are unable to consider the manuscript for publication in its present form.

The primary concern raised by both reviewers relates to the experimental design and the suitability of the biological material used. Specifically, the discovery that one of the ATAC-seq replicates was a hybrid, and that the stock labeled as D. albomicans may actually be D. nasuta, introduces significant uncertainty regarding the interpretation of species-specific chromatin accessibility. While we commend your transparency in acknowledging this issue in the supplementary material, it ultimately undermines the strength of the conclusions drawn.

Additionally, both reviewers have included additional aspects that can be considered to improve the clarity and strength of the work. We encourage you to consider these comments carefully should you decide to revise and resubmit your work. We appreciate your interest in the journal and thank you again for the opportunity to review your manuscript.

Referee #1 (Remarks to the Author):

The manuscript investigates the differences in chromatin accessibility in hybrids of Drosophila albomicans and D. nasuta species and their relationship with genetic differentiation. The authors suggest that chromosomal rearrangements can drive epigenomic divergence in hybrids, particularly in the Dot chromosome. They propose that the chromatin observed changes may play a role in reproductive isolation. This is an interesting topic given the limited data available, but the data presented do not sufficiently support the conclusions.

The major issue is that the stocks used are not the most appropriate material to address this study. The authors performed ATAC-seq on two representative strains with two biological replicates, but later discovered that one replicate was a hybrid, and that the stock considered to be D. albomicans seems to be D. nasuta. It is commendable that the authors acknowledge this problem and mention it in the supplementary material. However, in light of this, it is difficult to determine if the differences in chromatin accessibility are due to species differences, and whether they reflect functional or regulatory changes per se. Given these important shortcomings in experimental design, I cannot recommend the paper for publication in the journal.

I propose other aspects that the authors could address to improve the paper.

Some basic flaws in Material & Methods and through the manuscript could be addressed and justified:

-A clearer description of the stocks used, especially those used in ATAC-seq experiments, which is only included in the supplementary material. In the same way it will be very helpful to clarify that the so-called hybrids are introgressed lines and not crosses performed in the lab.

-More details on the ATAC-seq protocol and sequencing depth

-A brief explanation why some DAC regions differ between Deseq and EdgeR.

- Some figures lack necessary information. For example, in Figure 3B, the red dots mentioned in the legend are not visible in the figure. Similarly, in Supplementary Figures 10 and 11, the panels labeled A, B, C, etc., are not clearly identified.

- Provide a more detailed discussion based on the results obtained and relevant literature. Some paragraphs of discussion could be included in the introduction (e.g., lines 302-306). Conversely, there is little discussion about the genes present in the DAC regions and whether the high content of transposable elements in Dot chromosome could explain the observed differences.

- The conclusions should be stated more cautiously because they are not completely supported by the results.

Referee #2 (Remarks to the Author):

In this work, the authors investigated the modification of genomic regions of accessible chromatin (AC) as a consequence of chromosomal rearrangements (CRs). To explore this interesting and relatively under-explored topic, they selected two closely related Drosophila species with distinct sex chromosome constitutions that arose from chromosomal fusions, along with other minor rearrangements. Technically, the study is sound and employs the widely used ATAC-seq technique, a gold standard for obtaining information on AC. The analyzed data focused on AC in the testes of hybrids between D. albomicans and D. nasuta, and genome-wide fixation index (FST) estimates were used to identify regions with potential “accumulated” genomic differences and differences in AC. Together, these data were used to discuss how genomic and epigenetic architecture could contribute to the speciation process. Overall, the work is well presented. However, I have some comments regarding points that need clarification and aspects of data interpretation that could be addressed before possible consideration of the publication.

1 - The main data obtained was the identification of the Muller F chromosome as a hotspot of epigenomic and genetic divergence, despite its small size, is a noteworthy finding and may open new avenues of investigation. The introduction could be more directed to this aspect, than to directly to CRs. Or at least give a better panel of information for this element, it will orientate the reader about the impact of the work and of this element.

2 - Complementary RNA-seq data could strengthen the interpretation regarding the transcriptional consequences of the differences in accessibility.

3 - Although the data about Muller F is highlighted, other chromosomes or genomic regions would deserve a more detailed comparative analysis to reinforce the conclusion that initial changes tend to arise in heterochromatin/on Muller F.

4 - Some interpretations about the role of pioneer transcription factors still sound speculative. It would be important to support these hypotheses with direct evidence.

5 - The discussion could be more balanced, also providing examples of other models of chromosomal speciation to place the results in a broader evolutionary context.

6 - Considering that, as generally occurs in nature, the testis of Drosophila is a highly complex organ, the accessible chromatin (AC) could differ in distinct portions of this organ (pre-meiotic, meiotic, and post-meiotic cells). Why did the authors choose to analyze the whole organ?

7 - It is known that during spermatogenesis, the Muller F chromosome occupies a territory depleted of RNA polymerase II and is epigenetically silenced through processes such as hypermethylation and hypoacetylation, at least in Drosophila melanogaster. Consequently, this chromosome naturally exhibits a lower degree of transcription compared to the large autosomal elements. This information is relevant because the position of Muller F could influence its AC patterns, not directly due to the chromosome itself, but due to its position in primary spermatocytes. Some of these points could be better addressed in the discussion.

8 - Considering that epigenetic landscapes can differ dramatically between somatic and germline cells, the authors should justify the use of germline cells rather than somatic cells. Although this is briefly mentioned in the discussion, it should also appear in the Methods section to provide stronger motivation for selecting this organ compared to others.

9 - Why was the AC analysis focused primarily on the F element? The other chromosomes could be included, at least for comparative purposes.

10 - Based on Figure 1b, the X chromosome and F element show more uniform non-accessible chromatin, albeit with slight differences, and their overall accessibility is generally lower than that of autosomes (more evident to X chromosome). The text clearly indicates that the X chromosome has a lower degree of accessibility compared to autosomes. This pattern is consistent with X chromosome reduced expression during Drosophila (at least D. melanogaster primary spermatocytes) spermatogenesis and should be discussed. Additionally, if possible, it should be clarified whether differentially accessible chromatin (DAC) regions are associated with euchromatic or heterochromatic regions of the X chromosome, as these regions can exhibit distinct transcriptional and epigenetic profiles. In the discussion, the authors only address this superficially. Please also consider citing more recent literature on X chromosome inactivation in Drosophila (e.g., PMC7873209).

11 - Please clearly state that DAC values were normalized to chromosome size and gene content.

12 - In Results, topic 2, please clarify why the data from DESeq2 were selected for further analysis (provide the rationale).

13 - “Our study provides evidence for the existence of species that shows the transition of Muller F from sex-chromosome-like to autosome-like behavior in spermatogenesis.”. This pattern is not immediately clear, at least during my reading. Please clearly describe the supporting evidence in the discussion.

14 - In the discussion, the authors note that the study initially focused on the role of inversions in AC changes, but interesting patterns observed in Muller F prompted additional focus on this chromosome. However, in the introduction, the chromosomal rearrangement presented is a fusion. Please provide more information about chromosome inversions between the species in the introduction.

15 - In the conclusion, the authors state: “Our main objective was to understand how CRs and ACs affect the speciation in general and precisely how they shape the genome of hybrids of closely related species, D. albomicans and D. nasuta.”. However, the changes were much more evident in the Muller F, which does not directly contribute to karyotype differentiation between the species (e.g., large CRs observed in the emergence of the neo-sex system). Therefore, AC differences between the species may not be directly associated with CRs, but instead reflect ongoing changes specific to the F element. To clarify this, it would be important to study species with no CRs that are able to form hybrids. I suggest moderating the association between AC differences and CRs.

16 - It would be valuable to analyze AC differences between the two species, rather than in hybrid stocks. This would help determine whether the hybridization process influenced AC differences or if these differences are already established in natural populations and primarily reflect accumulated changes in Muller F.

17 - The title should be more focused to the main findings.
