## Supplementary material for "Chromatin accessibility differences between the hybrids of *nasuta-albomicans* complex of *Drosophila*": Review from journal 2

We have received two thorough expert reviews of your manuscript. Unfortunately, both reviewers agreed that the interpretation of your study is not reliable enough for publication at this time. Both reviewers felt that the changes necessary for publication are too immense to be done in the context of major revisions. However, both reviewers generously provided very helpful comments, which I hope you will find useful in continuing your work.

Reviewer(s)' Comments (if applicable):

Reviewer: 1

Comments to the Author

Summary:

The authors compare chromosomal accessibility between four strains of hybrids between D. albomicans and D. nasuta to explore how chromosomal rearrangement events have influenced regulatory differences between species. This is then linked to population-genetic data from the two species to try to infer the effects of selection on guiding these epigenetic differences.

Recommend explicitly defining “Robertsonian”

The lack of discussion of D. miranda’s neo-X and neo-Y chromosomes is strange.

Placement of lines 86-95 is a bit strange and breaks the flow of trying to explain CRs.

Line 92-95, the claim here is a bit unclear, so it’s difficult to assess to what extent this is true. It is a bit clearer on lines 107-109, but if this is the claim, it is also not necessarily true. There has been quite a long history of study of the relationship between chromosomal rearrangement/centric fusions with both regulatory changes (or lack thereof) and speciation.

https://doi.org/10.1038/s41559-022-01894-w

https://doi.org/10.1038/s41588-019-0462-3

https://doi.org/10.1073/pnas.0501847102

https://doi.org/10.1073/pnas.83.21.8245

Lines 96-97, please give readers an idea of the divergence times we are working with.

Lines 97-100, the precise chromosomal differences between species are critical for understanding the study design, but the lack of complete sentences makes it difficult to parse exactly what is happening.

Figure 1A, further illustration of the cartoon would be helpful in understanding the fusion that resulted in the neo-x and neo-y of D. nasuta and D. albomicans

Lines 101-103, again, the lack of even a basic phylogeny makes it incredible difficult to understand what is happening.

Line 111, what hybrids? These aren’t explained until the Methods section.

Lines 121-127 are very important and should be discussed more in the Introduction.

It’s unclear how these hybrids were found to have mostly D. albomicans or D. nausuta ancestry. This is critical information for interpreting all downstream analyses, and at the very least, the authors should make it clearer that this information is in the Supplementary Text. The most important piece of data, whether AAT1, AAT2, ANT1, or ANT2 have fused chromosomes or not is entirely missing.

It seems that AAT1 and AAT2 pairs and ANT1 and ANT2 pairs were used as biological replicates. This needs to be stated explicitly.

Figure 1B’s tracks are not labeled, so it is unclear how it is meant to be understood. There is some information in the caption, but it’s unclear how four strains were plotted here.

Lines 213-228 – what is the takeaway?

Lines 229-253 – please connect these results to the original questions of the manuscript

Lines 254-275 – this analysis can be motivated better.

Lines 276-292 – please guide readers through these results

Reviewer: 2

Comments to the Author

Overview:

To determine how chromosome rearrangements affect epigenetic states, Padma et al took advantage of the closely related species pair D. albomicans and D. nasuta to determine whether hybrids have altered epigenetic states. The former species had two recent Robertsonian fusions of the autosomes to the X and Y, creating young neo-sex chromosomes that are absent in the latter. The authors claim to have found, on the dot chromosome that widely differ in chromosome accessibility between two hybrid strains and claimed that the region overlaps with an island of elevated Fst.

Comments:

I have several major concerns about this manuscript and can not endorse its publication. The main concerns include the lack of congruence between motivation and results, the mysterious origin or the “hybrids”, the lack of investigation into the pure species, and overclaim of elevated Fst and the relevance to speciation, dubious association with pioneer factors.

“Chromosome rearrangements” and differential accessibility are unrelated

The authors motivated study in trying to understand how rearrangements can affect changes in epigenetic states. The two species differ in the presence of a pair of Robertsonian fusions. First, lumping this Robertsonian fusion in with other types of chromosomal rearrangements is somewhat misleading because the fusion itself does not prevent recombination between the fused and unfused versions (See Wang et al 2022) , unlike other rearrangements like inversion. Second there are known inversions between the the two species (see Mai and Bachtrog 2021, https://doi.org/10.1101/2021.06.01.446624) that the authors do not mention. So even if any relationship can be claimed between chromatin state and chromosome rearrangements, they cannot differentiate between different types of structural variation. The even bigger issue is that the Robertsonian fusion of the neo-sexes is actually irrelevant to their main results which pertain to Muller F, a completely separate chromosome.

The “hybrids” are of unknown origin and genotypes

The authors collected ATAC-seq on hybrids that appear to have different extent of nasuta vs albomicans ancestry. As far as I can tell, these hybrids were mislabeled or contaminated stocks that were identified to be more similar to one or the other species, which the authors realized after the ATAC-seq data had been collected. The authors go into extensive length in the supplementary data to show that these hybrids have introgressed genotypes, but the important question is whether and how these genotypes actually affect the ATAC-seq results. Are these stock inbred or are they contaminated lines with segregating heterozygosity? To make sense of the accessibility data, the authors need to first figure out what the genotype of Muller F is between the “hybrid” samples. Using inbred strains of the two species are obviously better choices.

Island of elevated Fst and implication to speciation

The elevated level of Fst on the dot chromosome is an interesting result but I am not entirely sure what to make of it. Muller F in Drosophila famously does not recombine, and one of the consequences of this is that the entire chromosome should have similar rates of divergence, due to linked selection/drift. Island of high Fst, which usually represents regions that do not recombine between populations, are islands because neighboring regions are able to recombine which therefore encourages gene flow. In this instance the entire chromosome cannot recombine, yet we still see regions with drastically different Fst. I wonder whether the pattern is driven by increased repeat content leading to more “blacked out” windows which may then lead to overestimates of differentiated positions (perhaps the same may also be occurring with the same region showing the most DAC). It is formally possible also that part of the chromosome has recombined - perhaps not out of realms of possibility given that these species have been suspected to have male recombination. Again, this is an interesting result that deserves much more attention. The link to “speciation”, “reproductive isolation”, reproductive barrier seems completely unfounded though. These species have no reproductive isolation as they are able to make viable offsprings. The only suggestion of hybrid incompatibility appear to arise as recessive incompatibilities in 2nd, 3rd generation hybrids in the form of sterility and reduced fertility.

In the DAC, the authors claimed to have found “higher binding affinity” (whatever that means) to several transcription factors including the pioneer factors zld, Trl and Clamp. First, it is not clear whether these pioneer factors even have the same recognition motif in the species as D. melanogaster where they are characterized. Second, the pioneer factor activity of these genes are supposed to occur during early embryogenesis, so it is not clear what their role in the germline could be.
