## Supplementary Figures for "Chromatin accessibility differences between the hybrids of *nasuta-albomicans* complex of *Drosophila*"

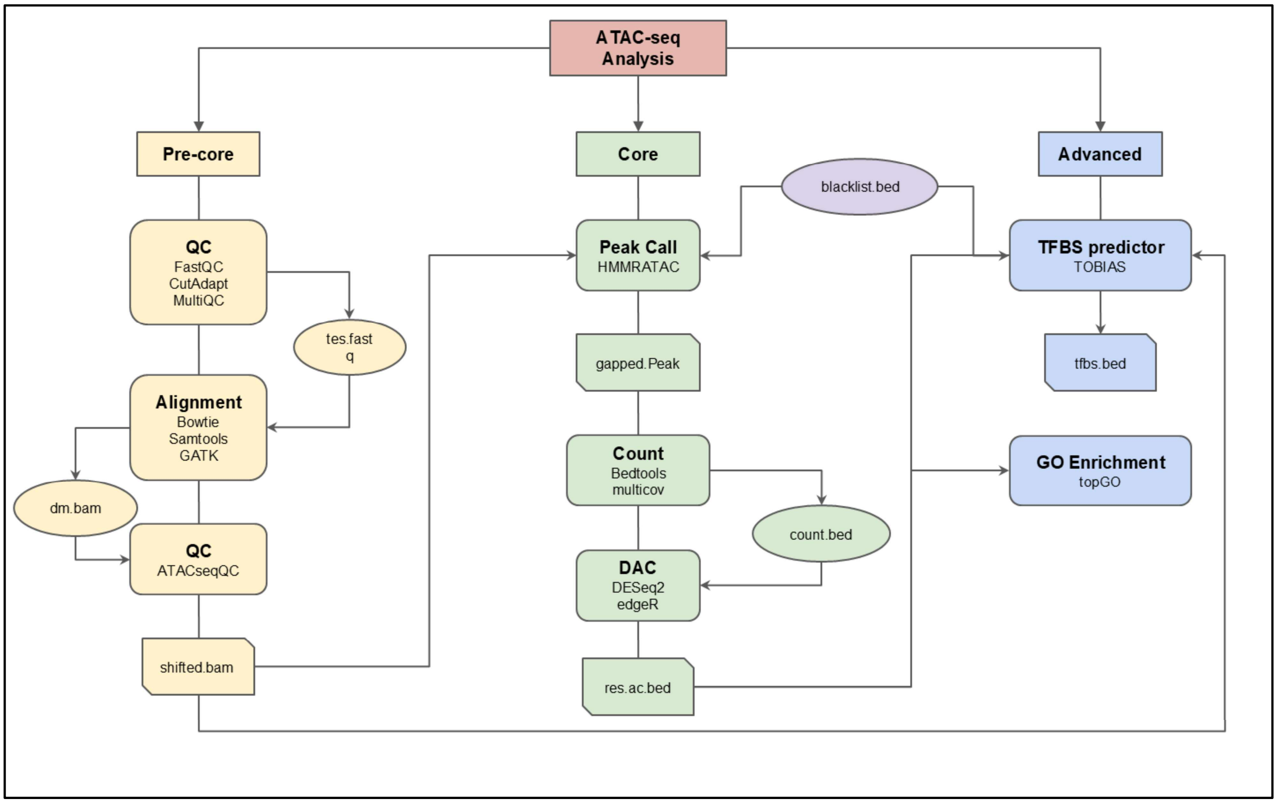


**Supplemental_Fig_S1: Outline of ATAC-seq analysis**

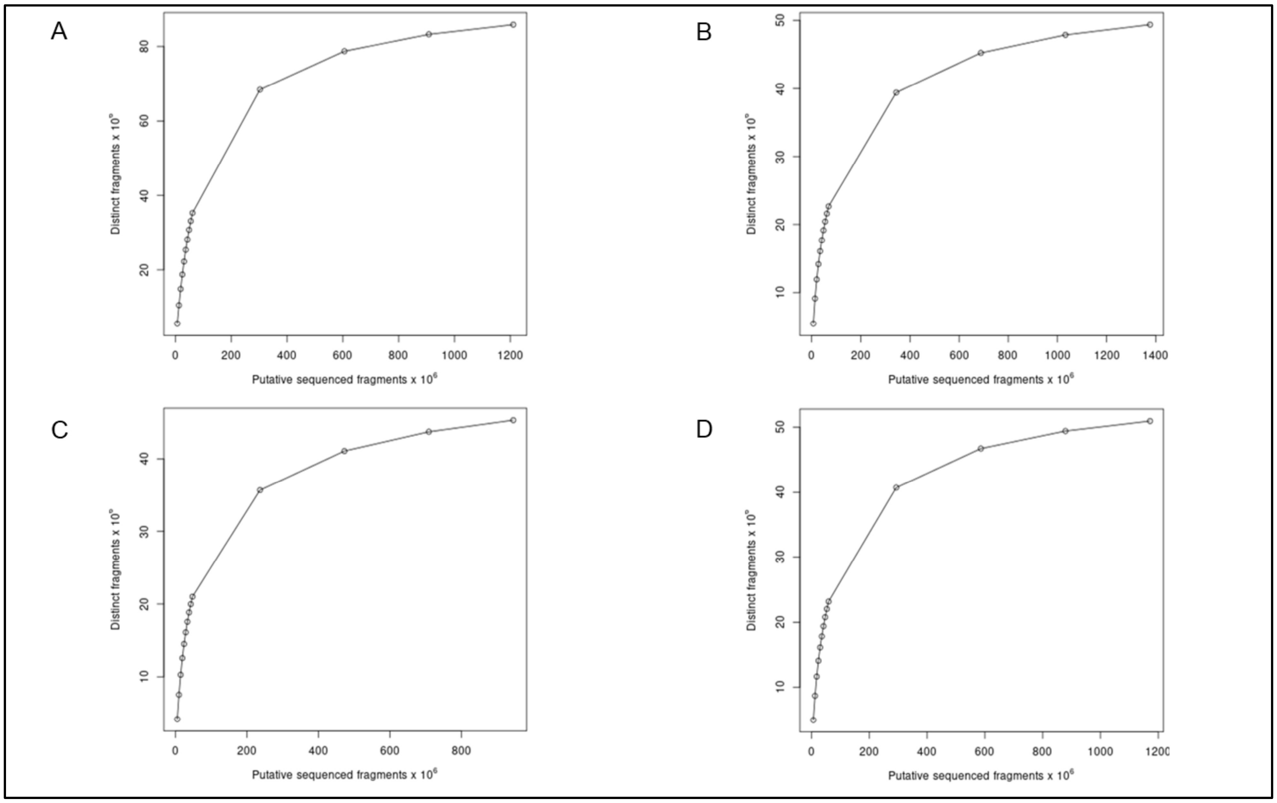


**Supplemental_Fig_S2: Estimation of ATAC-seq library complexity**

**A** and **B** Hybrid identifying with *D. albomicans* replicates, **C** and **D** Hybrid identifying with *D. nasuta* replicates

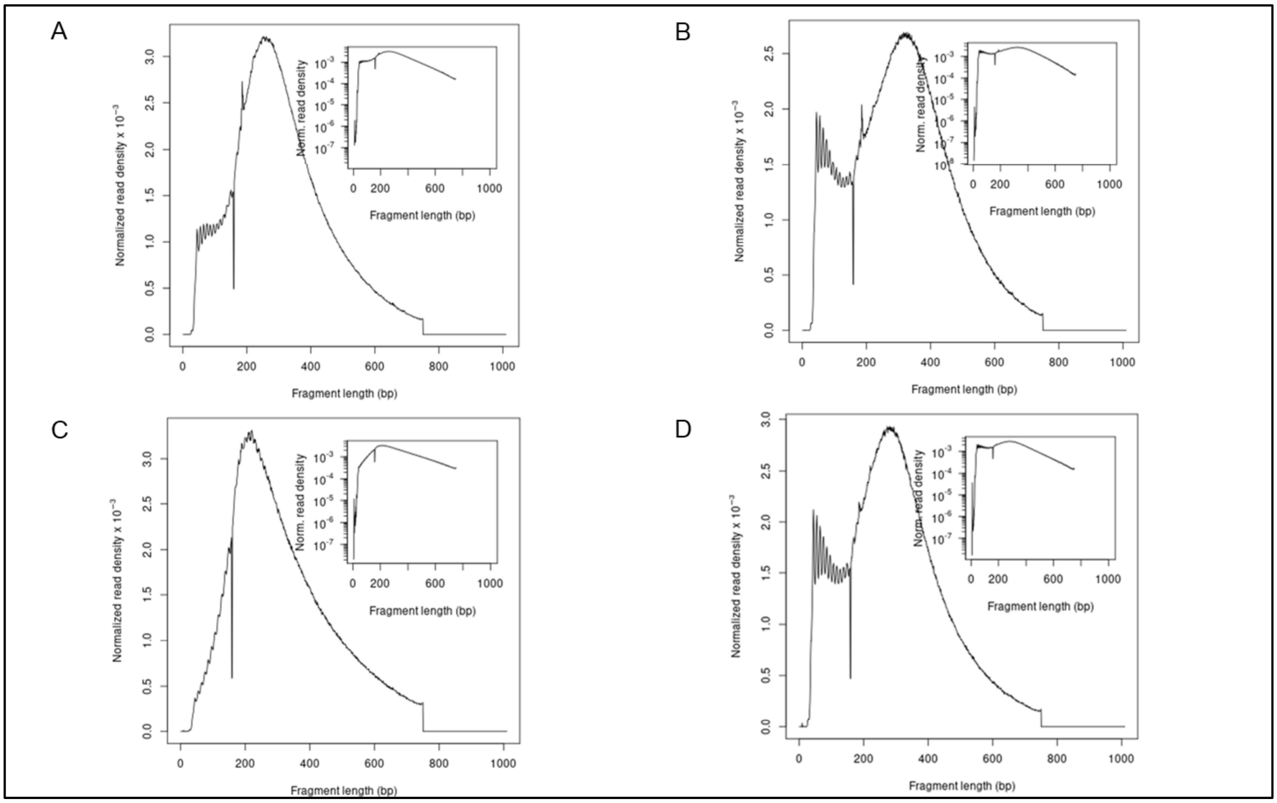


**Supplemental_Fig_S3: Fragment length distribution in aligned reads**

**A** and **B** Hybrid identifying with *D. albomicans* replicates **C** and **D** Hybrid identifying with *D. nasuta* replicates

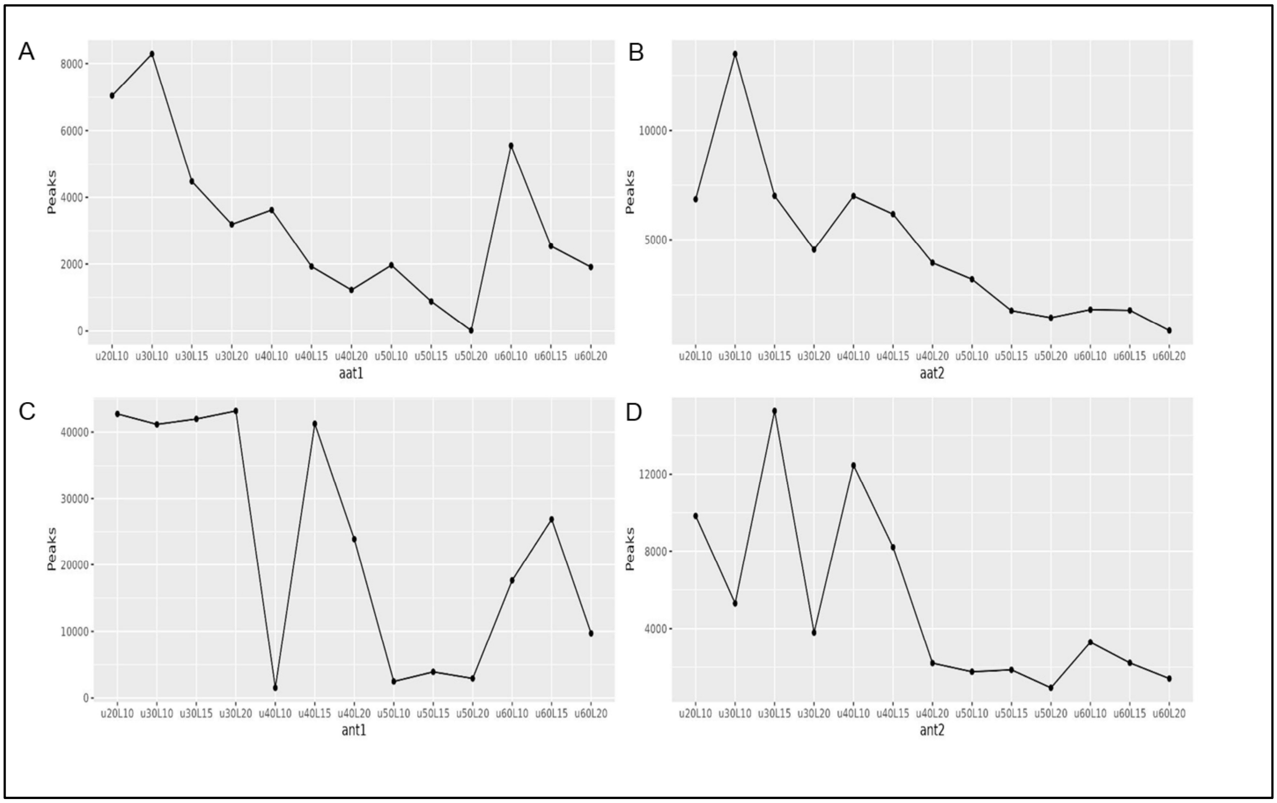


**Supplemental_Fig_S4: Number of peaks across models in ATAC-seq datasets**

**A** to **D** peaks of 13 models in four samples. We have selected the models which produced the highest number of peaks (sensitive) for the downstream analysis in our study.

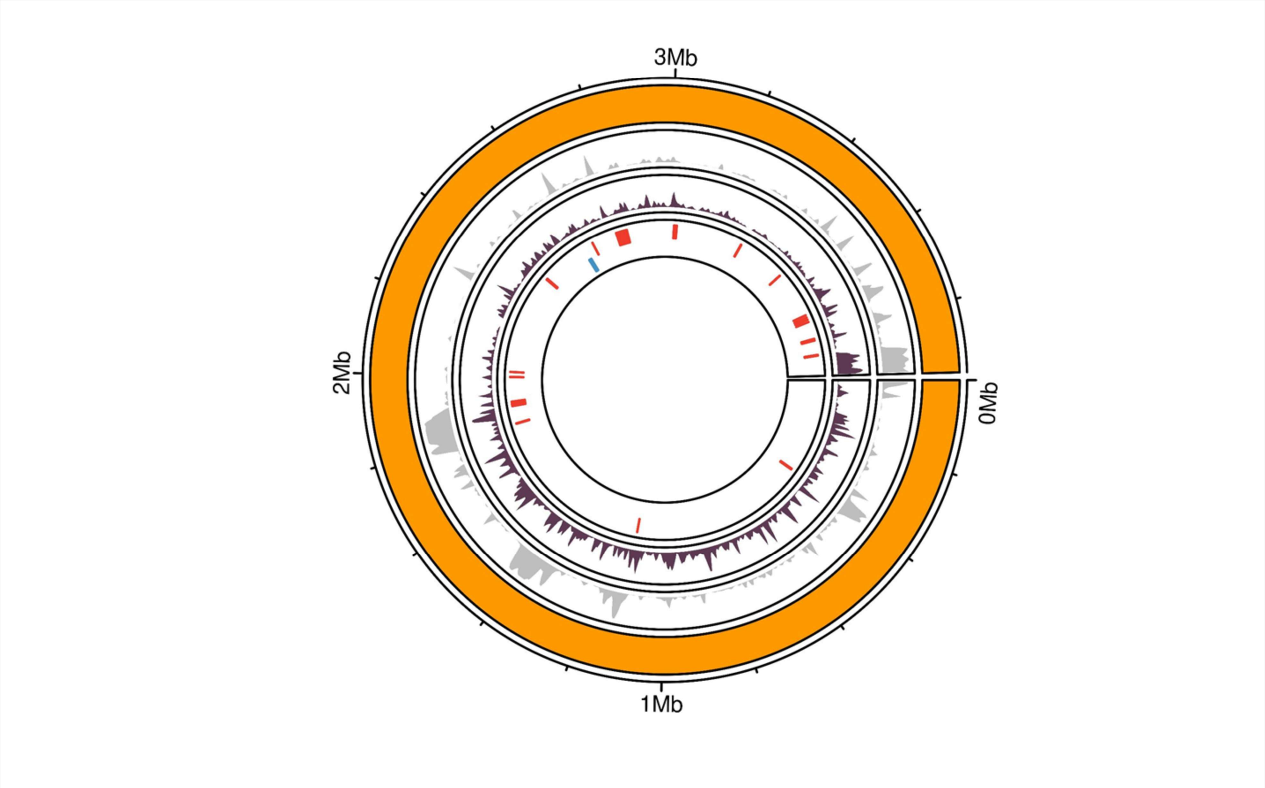


**Supplemental_Fig_S5: Accessible chromatin (ACs) of Muller F/chromosome 4**

Accessible chromatin (ACs) of testes in hybrids identifying with *D. albomicans* and *D. nasuta,* with Muller element color, the outermost and second outermost tracks represent 4th chromosome, and non-accessible chromatin density, respectively. The innermost track represents differential accessibility (DACs), blue and red bars represent high and low accessibility regions in hybrid identifying with *D. albomicans*. The second innermost track represents the transposable element density.

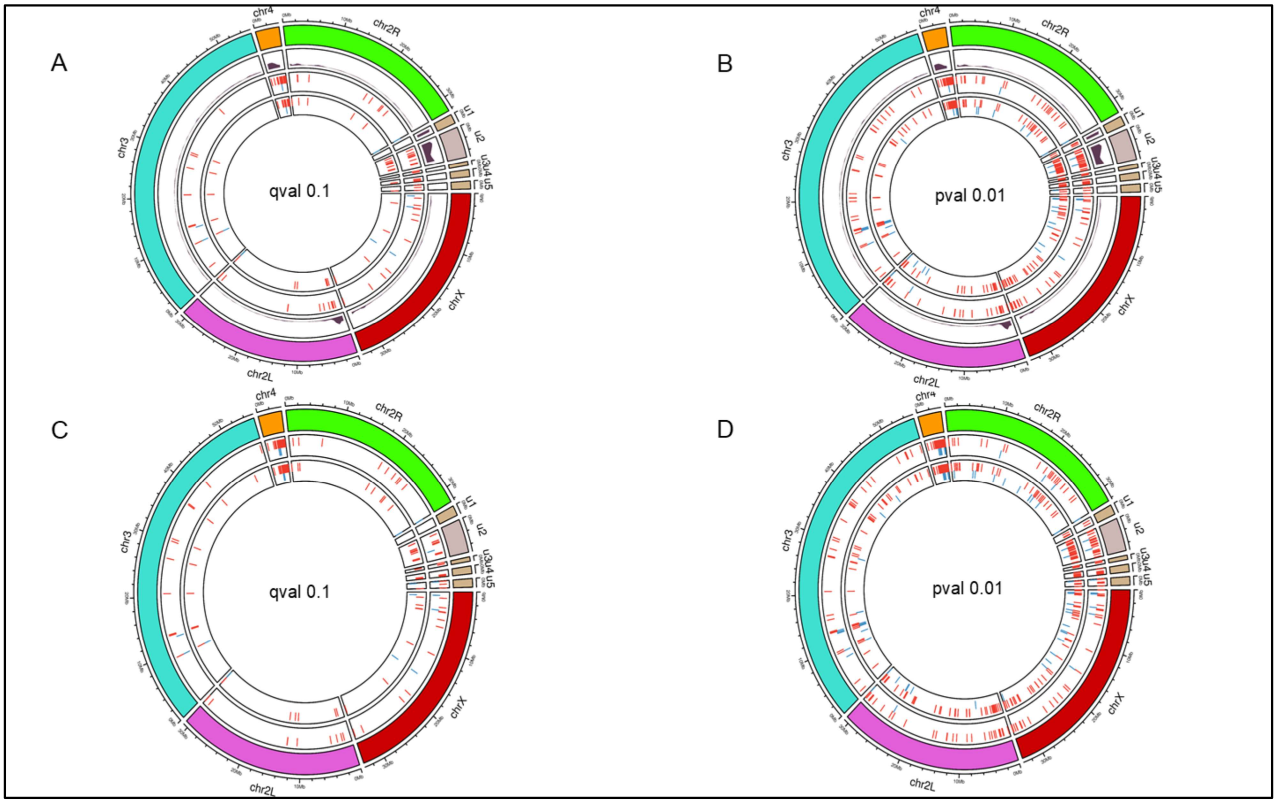


**Supplemental_Fig_S6: Two methods analyzing DACs and annotated transposon-free regions of ACs in hybrid identifying with *D. albomicans***

**A** and **B** represent the DACs of padj-value 0.1 and p-value 0.01, respectively. The outermost and second outermost tracks represent the chromosomes and transposons found in the genome, respectively, whereas innermost and second innermost represent the edgeR and DESeq2 found DACs. **C** and **D** represent the DACs without annotated transposons of padj-value 0.1 and p-value 0.01, respectively. The outermost track represents the chromosomes, and the innermost and second innermost represent the edgeR and DESeq2 found DACs, respectively.

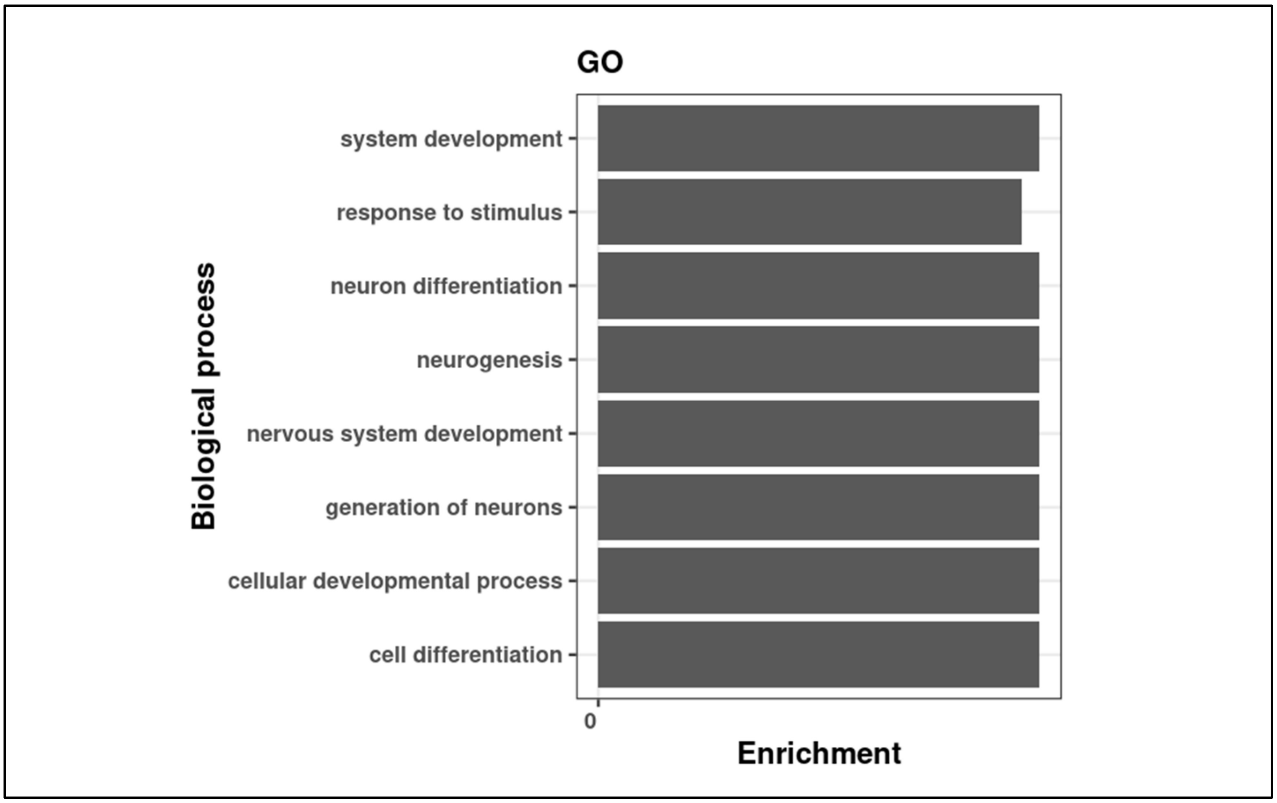


**Supplemental_Fig_S7: GO enrichment of testes in** hybrids identifying with ***D. albomicans*** **and** ***D. nasuta***

The GO enrichment analysis using topGO, R package. Based on Kolmogorov-Smirnov test in classic method.

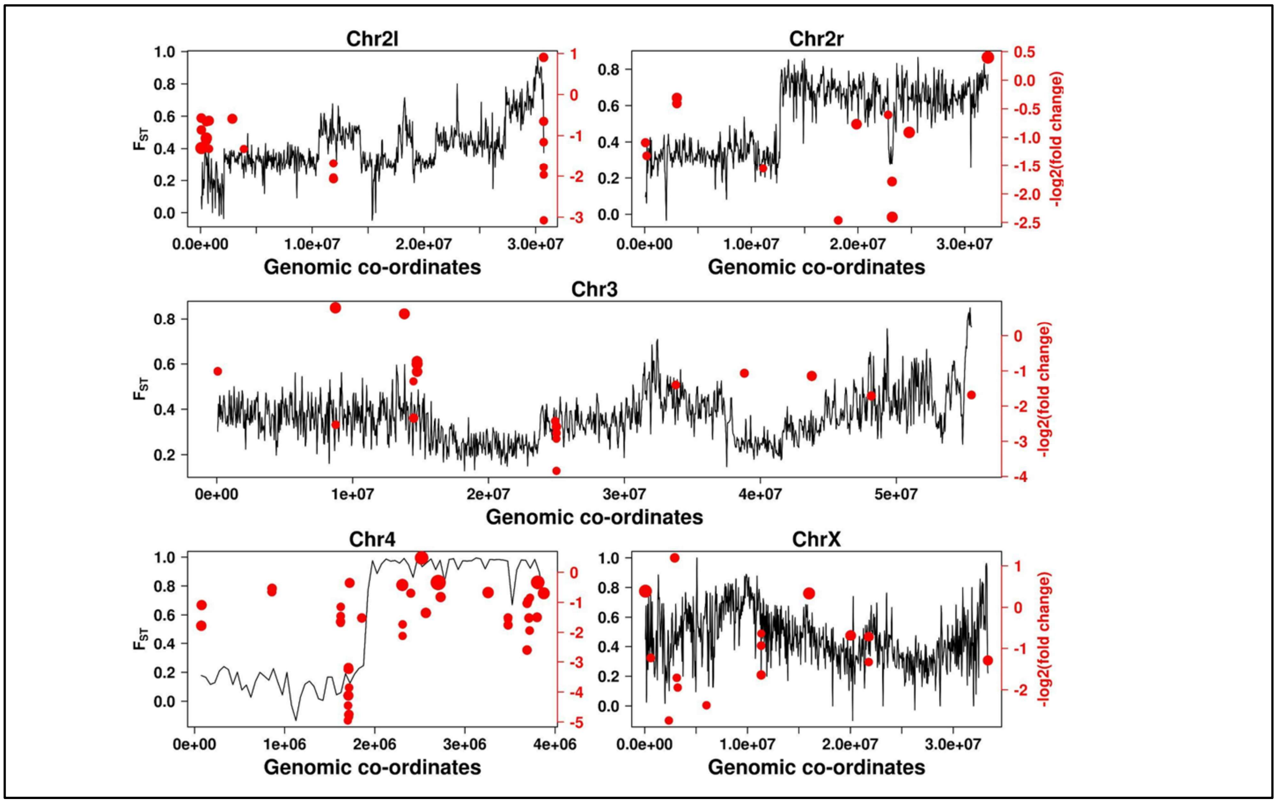


**Supplemental_Fig_S8: Genomic landscape of differentiation with regions of significant differential accessibility for *D. albomicans* (IND) vs *D. nasuta* (IND) population pair.**

$F_{ST}$ calculated for 50 Kb non-overlapping windows for each chromosome is plotted on the Y- axis (black). On the second Y-axis (red), -log2(fold change) of significant differential accessibility regions plotted with dot size proportional to length of the region. Due to greater range of lengths of regions, the dot sizes were log10 normalised. On chromosome 4 numerous bigger differential accessible regions can be observed overlapping with high Fst regions.

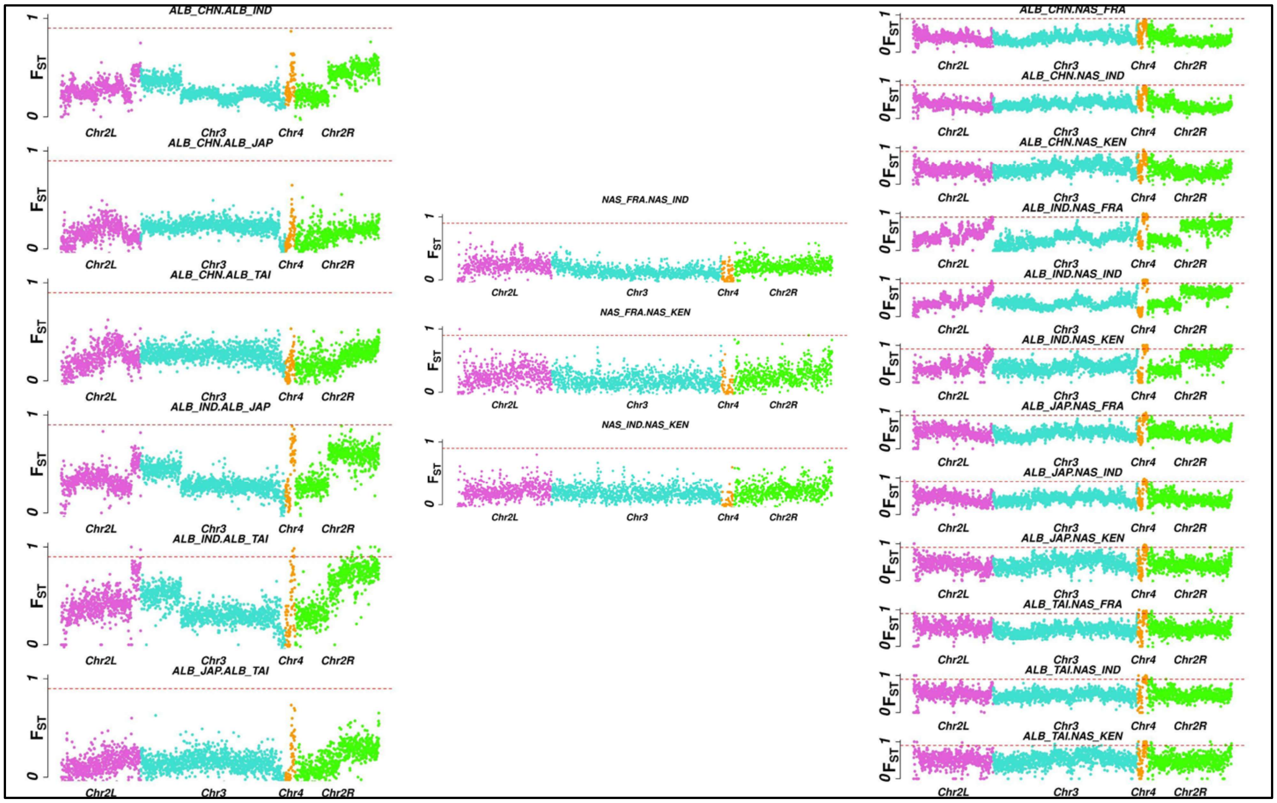


**Supplemental_Fig_S9: Genomic landscape of differentiation in *D. albomicans***
Pairwise genetic differentiation (FST) in 50-kb non-overlapping windows across the genome between *D. albomicans* population pairs. Colours represent respective muller elements; horizontal dotted red line demarcates FST estimate of 0.9. JAP and TAI population show higher differentiation against IND population, which suggests possible existing population structure among these pairs.

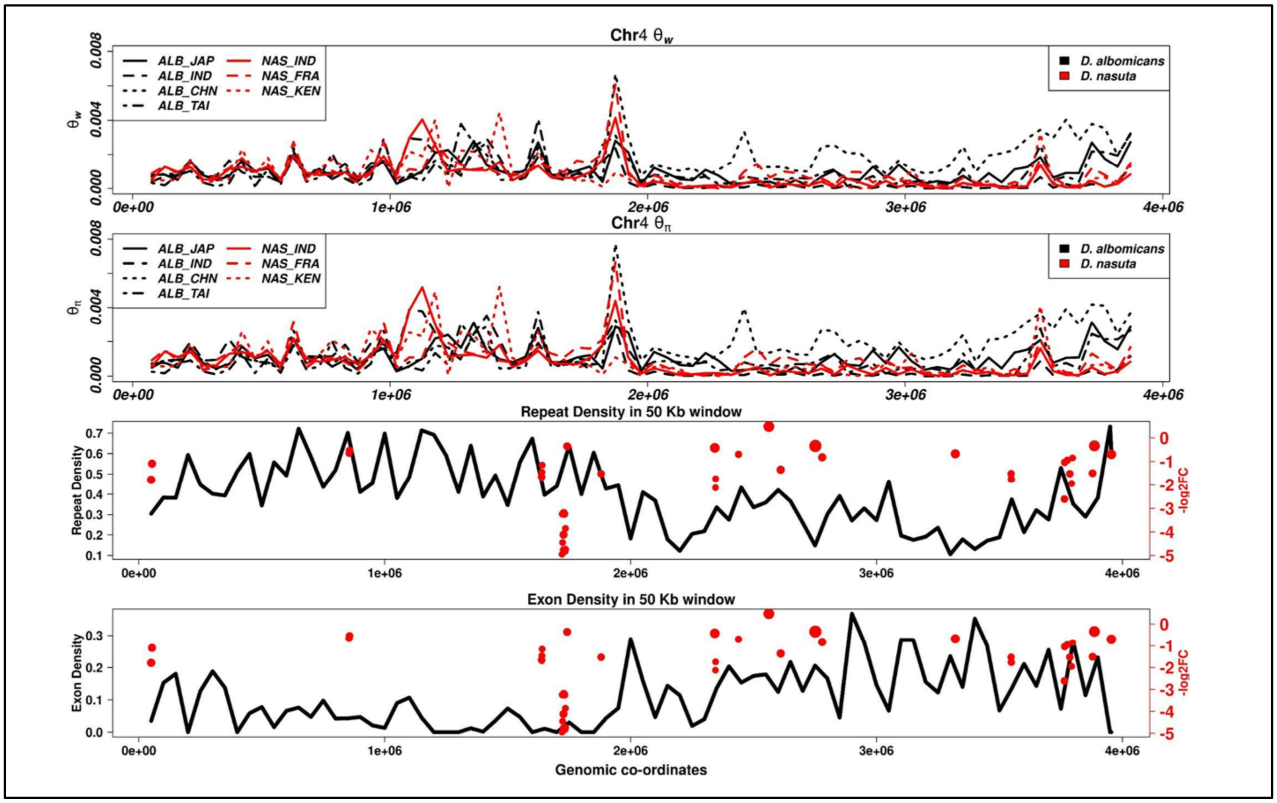


**Supplemental_Fig_S10: Genomic characteristics of Chromosome 4**

A) Distribution of nucleotide diversity (Watterson's theta) in 50 Kb non-overlapping windows across chromosome 4 for all the populations. Overall diversity pattern is conserved except CHN population which seems to harbour higher nucleotide diversity compared to other populations.

B) Distribution of pairwise nucleotide diversity (Theta pi) in 50 Kb non-overlapping windows across chromosome 4 for all the populations.

C) Density of repetitive elements in 50 Kb non-overlapping windows across chromosome 4 on first Y-axis (black). On the second Y-axis (red), -log2(fold change) of significant differential accessibility regions is plotted with dot size proportional to length of the region. Due to greater range of lengths of regions, the dot sizes were log10 normalised.

D) Density of exons 50 Kb non-overlapping windows across chromosome 4 on first Y-axis (black). On the second Y-axis (red), -log2(fold change) of significant differential accessibility regions is plotted with dot size proportional to length of the region. Exon density is greater in the later half of the chromosome.

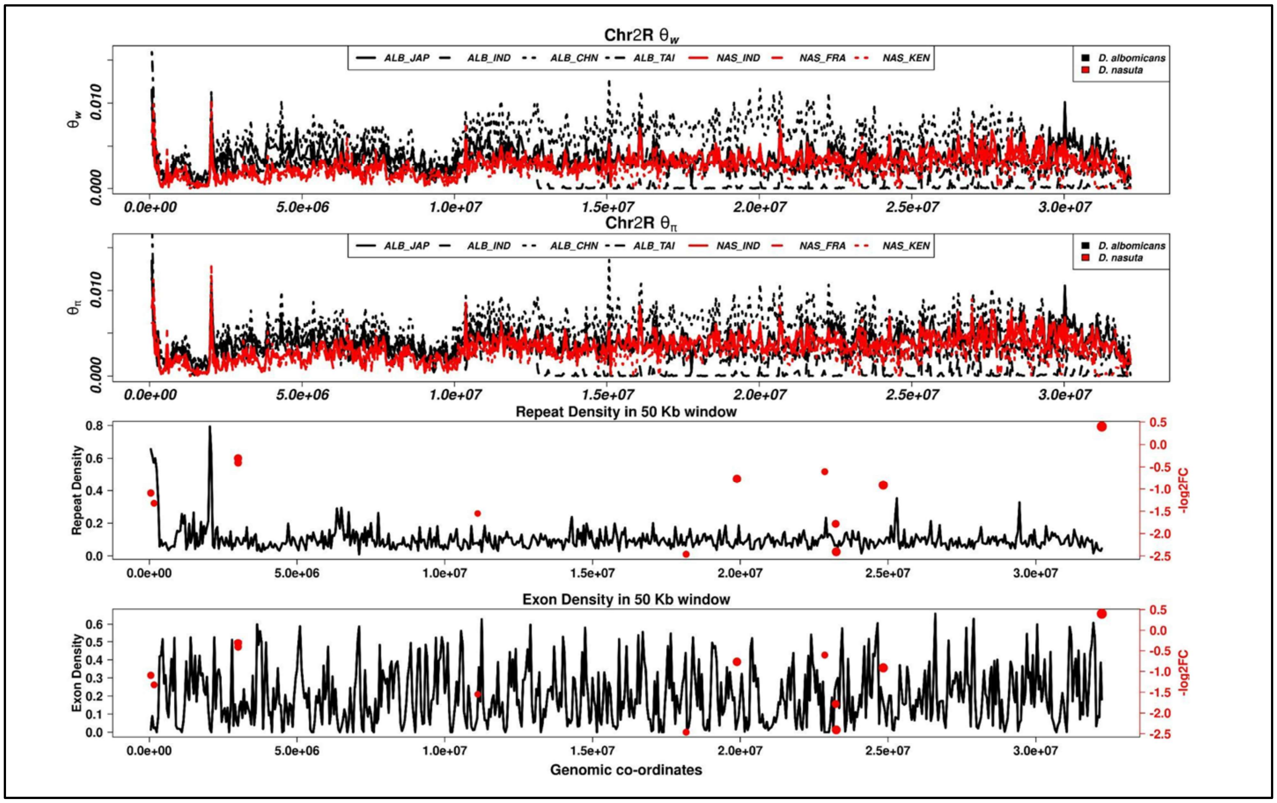


**Supplemental_Fig_S11: Genomic characteristics of Chromosome 2R**

A) Distribution of nucleotide diversity (Watterson's theta) in 50 Kb non-overlapping windows across chromosome 2R for all the populations. Overall diversity pattern is conserved except CHN population which seems to harbour higher nucleotide diversity compared to other populations.

B) Distribution of pairwise nucleotide diversity (Theta pi) in 50 Kb non-overlapping windows across chromosome 2R for all the populations.

C) Density of repetitive elements in 50 Kb non-overlapping windows across chromosome 2R on first Y-axis (black). On the second Y-axis (red), -log2(fold change) of significant differential accessibility regions is plotted with dot size proportional to length of the region. Due to greater range of lengths of regions, the dot sizes were log10 normalised.

D) Density of exons 50 Kb non-overlapping windows across chromosome 2R on first Y-axis (black). On the second Y-axis (red), -log2(fold change) of significant differential accessibility regions is plotted with dot size proportional to length of the region. Exon density is does not have any pattern like Chromosome 4.
