## Supplementary Text for "Chromatin accessibility differences between the hybrids of *nasuta-albomicans* complex of *Drosophila*"

**Identification and quantification of hybrid ancestry in sampled individuals**

Initially, *D. albomicans* (Okinawa strain, Texas collection, USA, 3045.11) and *D. nasuta* (Coorg, India 201.009) were collected from the Drosophila Stock Center, University of Mysore, India, for assessing the differential chromatin accessibility landscape between the two species using ATAC-seq. The BAM files of the two ATAC-seq samples (with two replicates each) and those of the individual used for population genetic analysis **(Supplemental Table S2)** were analysed together using ANGSD version: 0.935-53-gf475f10 (Korneliussen et al. 2014) to call genotypes with stringent filtering criteria (-SNP_pval 2e-6 -minMapQ 30 -minQ 20 -minMaf 0.05 -uniqueOnly 1 -remove_bads 1 -only_proper_pairs 1 -trim 0 -C 50 -baq 1 -setMinDepthInd 10) to maintain the high quality of the data. The called genotypes were then input for NgsAdmix (Skotte et al. 2013) to estimate individual admixture proportions.

In the admixture plots, each vertical bar on the x-axis represents a single individual included in the analysis, grouped and often sorted by population or sampling location for visual clarity. The y-axis indicates the proportion of that individual's genome derived from each inferred ancestral population (ranging from 0 to 1). Each colour in the plot corresponds to a different ancestral population cluster inferred by the NgsAdmix algorithm at a given value of K, representing the assumed number of ancestral populations. For example, at K=3, three distinct colours indicate the contributions from three genetic clusters. The height of each colour segment within a bar reflects the fraction of the individual's genome associated with that particular cluster. Individuals with multiple colours are admixed, meaning they have ancestry from more than one genetic background. In contrast, individuals with a single dominant colour are inferred to have ancestry primarily from a single population. This visualisation facilitates the identification of hybrids, population structure, and potential misassignments of species identity.

The individuals included in the admixture analysis from left to right are AAT1 and AAT2 (ATAC-seq samples from *D. albomicans* lab stock), *D. albomicans* natural populations (with three individuals each) from China, India, Japan, the lab population ascribed to *D. albomicans,* followed by *D. albomicans* natural population from Taiwan. The next samples are ANT1 and ANT2 (ATAC-seq samples from *D. nasuta* lab stock), followed by *D. nasuta* natural populations (with three individuals each) from France, India, and Kenya. The final three individuals are the lab population ascribed to *D. nasuta*. The same order is followed in all plots.

The admixture proportions at K of 2 (see below figure) distinguish the *D. albomicans* and *D. nasuta* samples obtained from different countries (China, India, Japan and Taiwan vs France, India and Kenya). Interestingly, it was noted that the two individuals (AAT1 and AAT2) from the *D. albomicans* stock have an ancestry matching *D. nasuta.* Similarly, the two individuals (ANT1 and ANT2) from the *D. nasuta* stock have an ancestry matching *D. nasuta* in the case of one individual and a hybrid ancestry (multiple colours in the same bar) in the other.

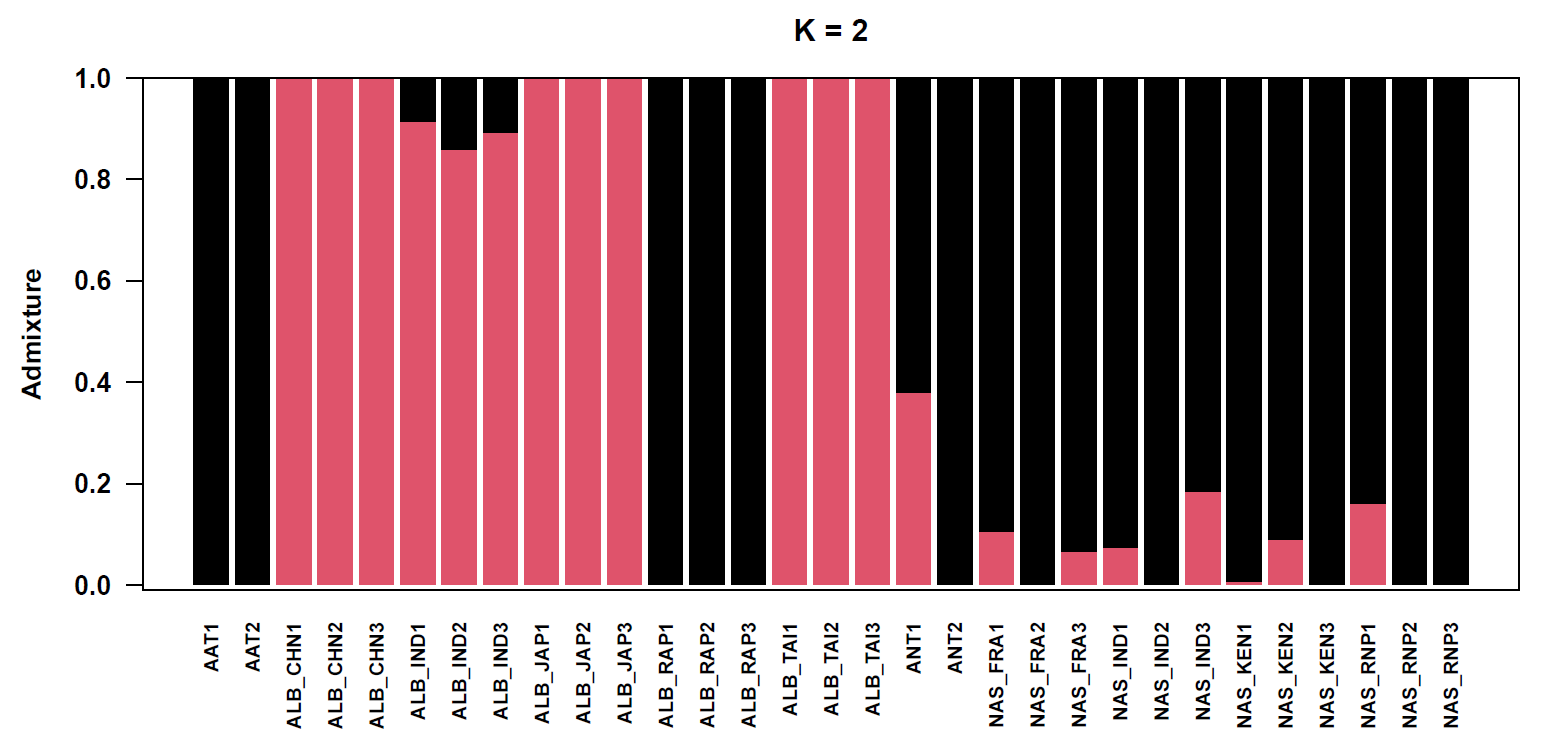

The estimates at k=3, identify sub-structure within *D. albomicans* populations. Specifically, the *D. albomicans* population from India was found to have *D. nasuta* ancestry. Another important pattern was the clear separation of the lab and natural populations. The ANT1 sample continued to have mixed ancestry, similar to the pattern seen at K=2.

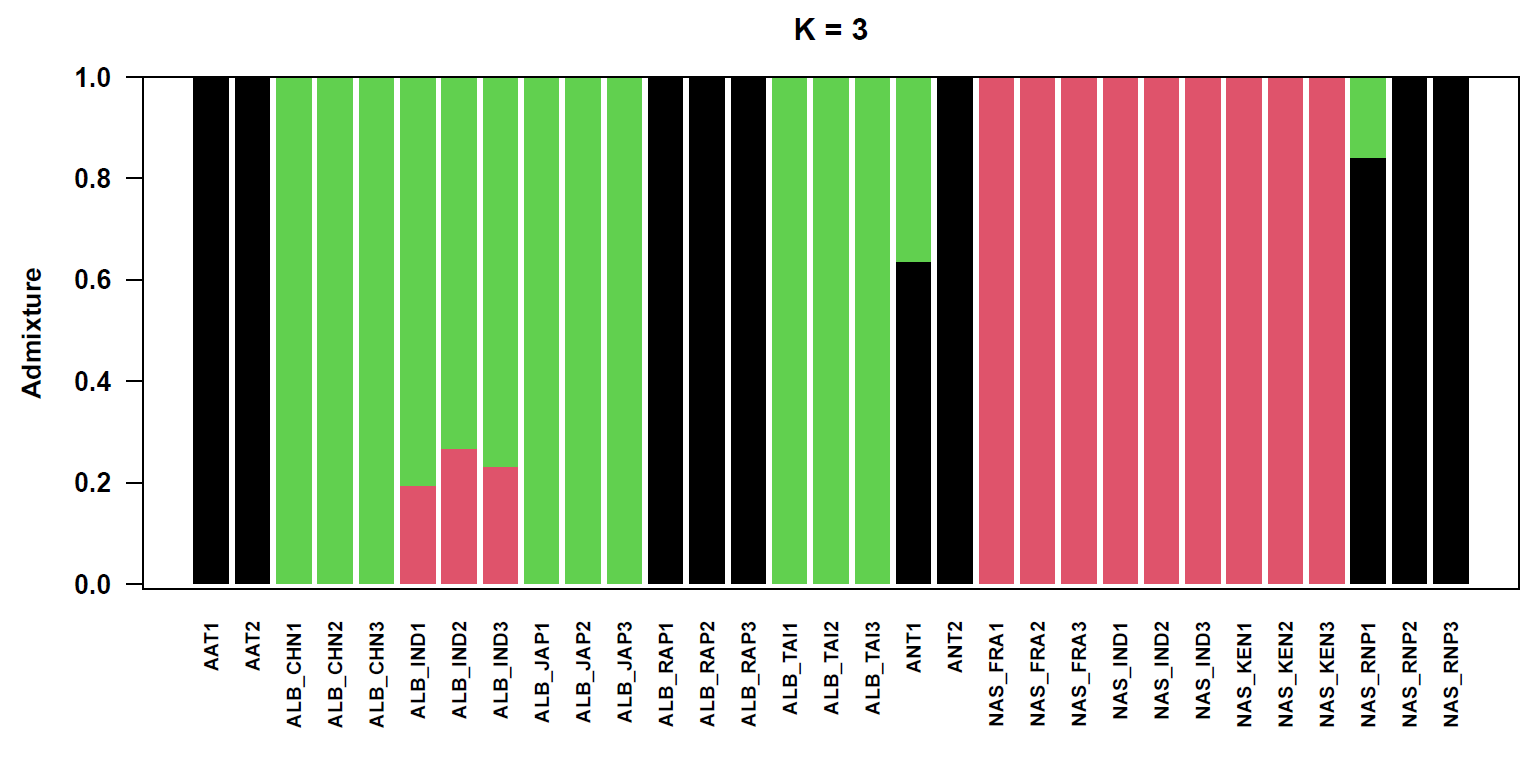

At a K of 4, the separation between lab and natural populations continued. Additional sub-structure was noticed within *D. albomicans* populations in contrast to the lack of sub-structure in *D. nasuta*. Specifically, the *D. albomicans* population from China was intermediate between the population from India and the populations from Japan and Taiwan.

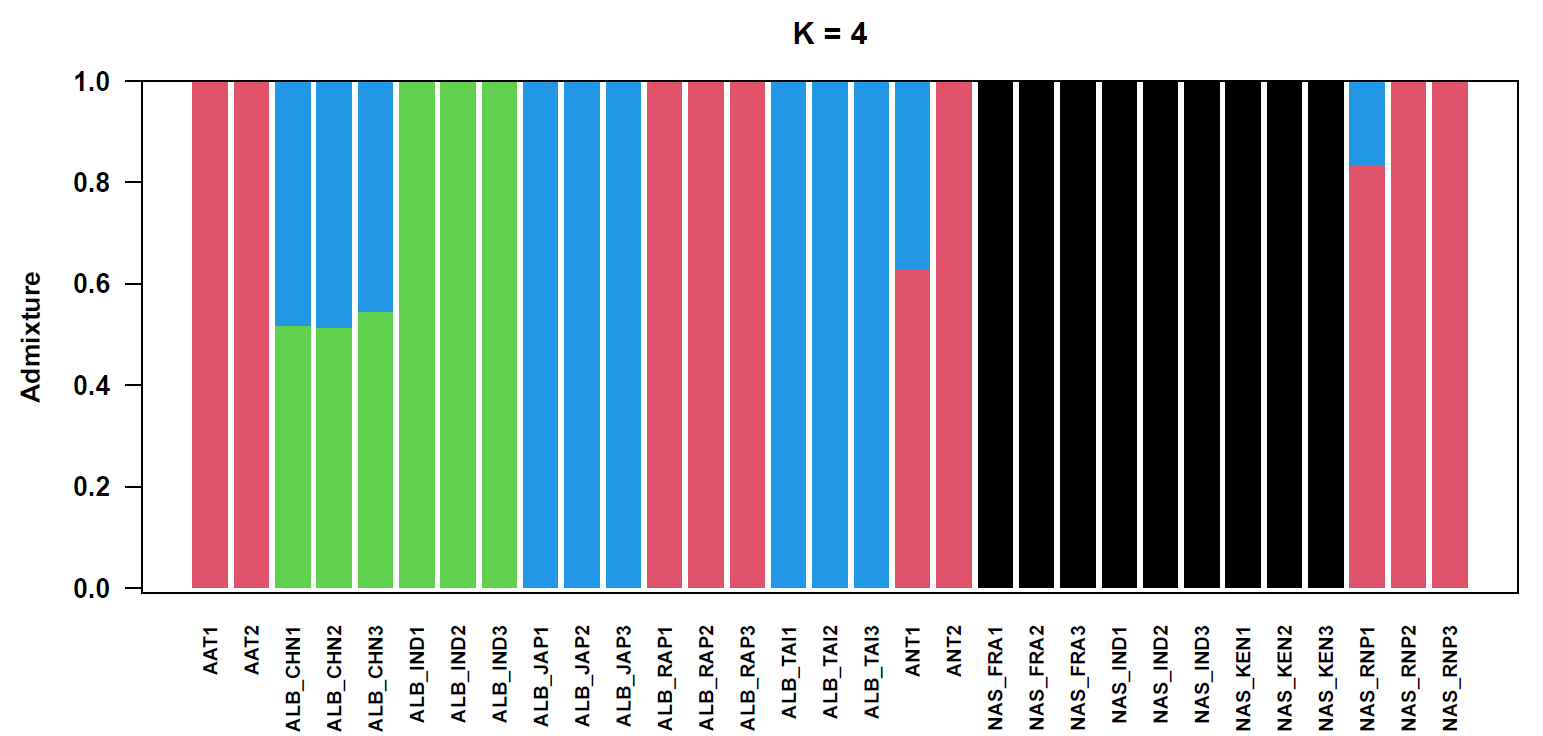

At higher values of K (i.e., 5 and 6), additional sub-structure was evident in *D. albomicans* populations. Moreover, the ANT1 and ANT2 individuals appear as hybrids, while the AAT1 and AAT2 are very distinctive.

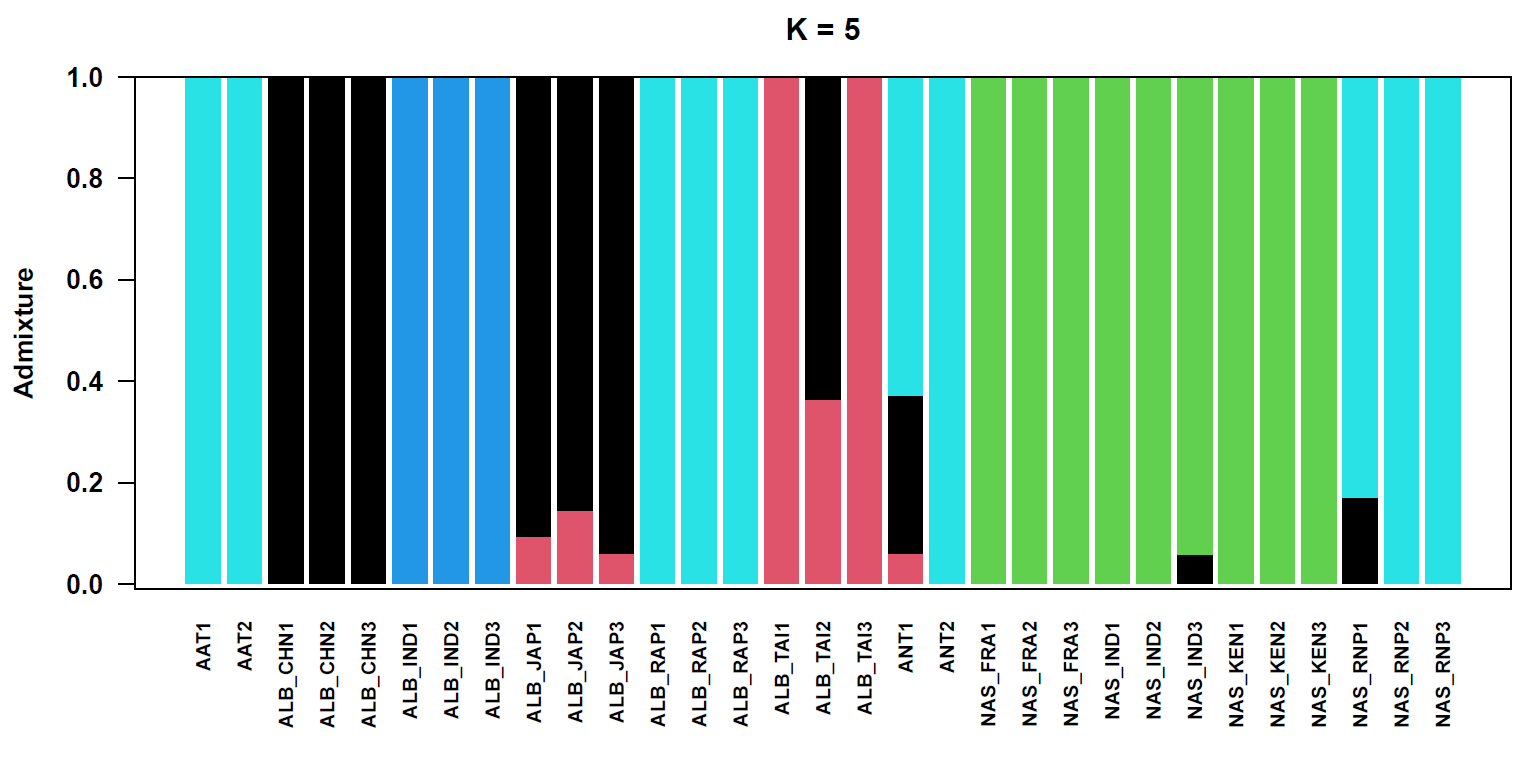

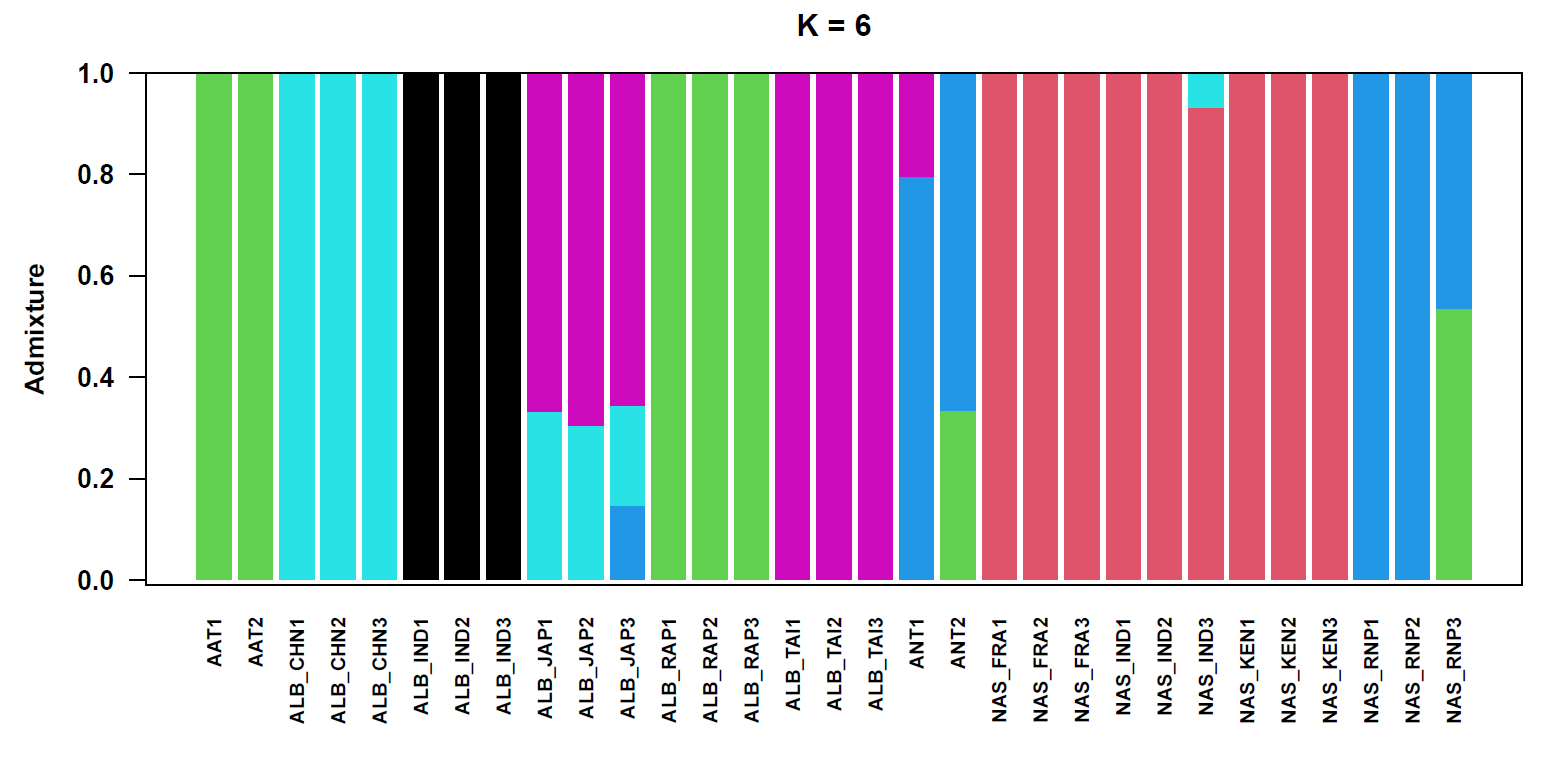

At a K of 7, the sub-structure within *D. nasuta* populations can be seen.

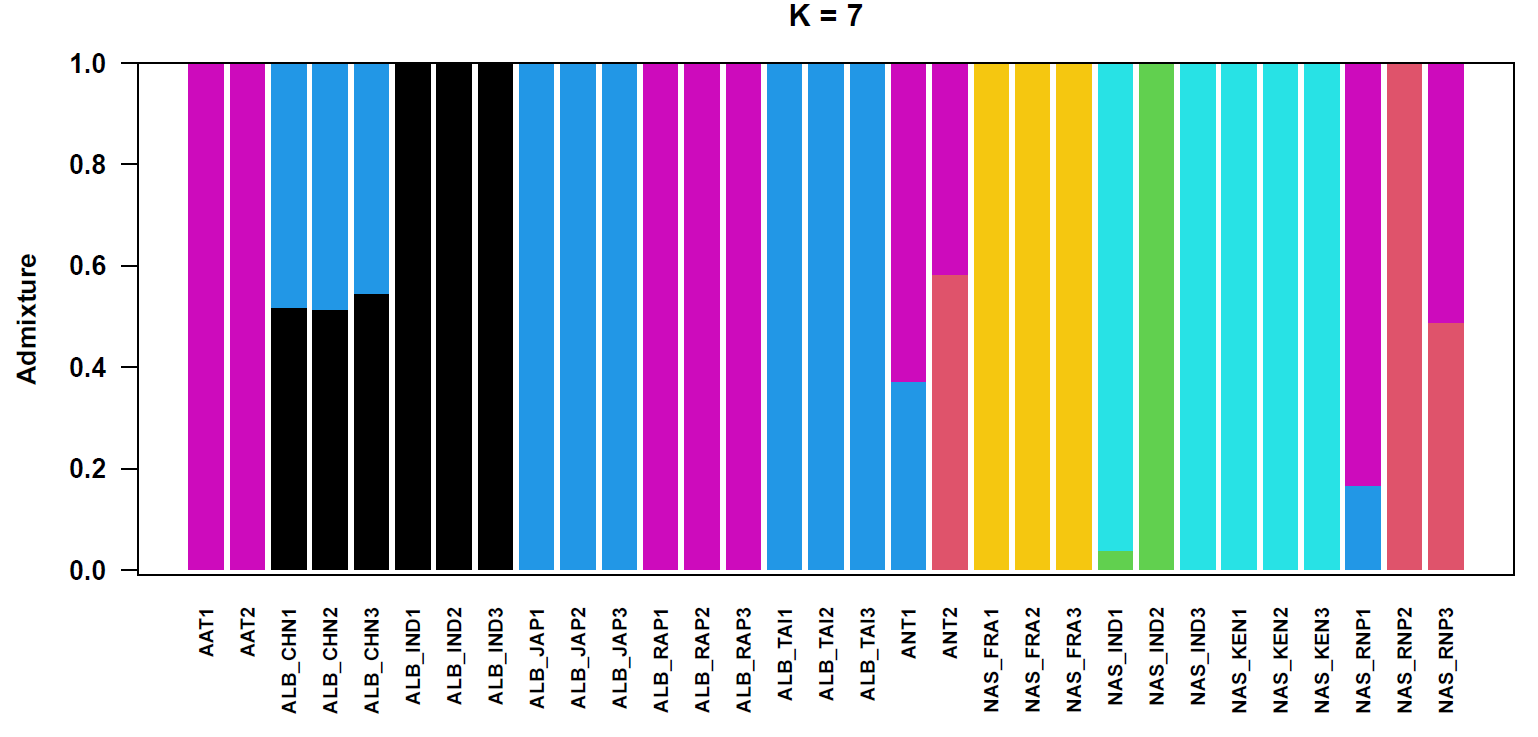

The admixture plots presented provide a comprehensive visual summary of the genetic ancestry of individual samples at increasing levels of population resolution (K = 2 to K = 7). At K=2, a clear bifurcation between *D. albomicans* and *D. nasuta* is observed, consistent with species-level differentiation. However, several individuals (notably AAT1, AAT2, ANT1, and ANT2) show ancestry components inconsistent with their reported species designation, revealing instances of cryptic hybridisation. At higher K values, additional population structure emerges, particularly within *D. albomicans*, indicating geographically correlated genetic subgroups. The ANT1 individual maintains a hybrid ancestry profile across all K levels, suggesting it is a first- or second-generation hybrid. In contrast, AAT1 and AAT2, although derived from a *D. albomicans* stock, consistently group with *D. nasuta* populations, suggesting either sample mislabeling or ancestral introgression. The admixture profiles thus highlight substantial cryptic genetic complexity within laboratory stocks previously considered pure species.

Based on the results of the admixture analysis, the sampled individuals from the stock appear to have hybrid/*D. nasuta* ancestry as assessed by the public population genomic data. We used principal component analysis (PCA) to evaluate the sampled individuals' genetic composition further.

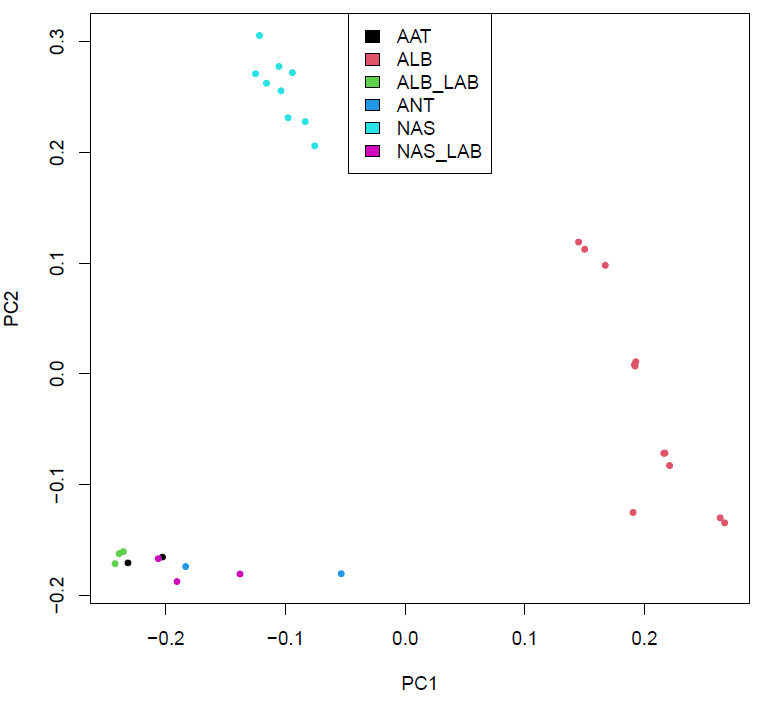

The PCA results clarify the drastic difference between the lab and natural populations. The clustering in *D. nasuta* is much more compact than *D. albomicans* and mirrors the pattern seen in the admixture plots. Overall, our analysis of the genetic composition of the samples from the stock centre compared to the natural populations sampled suggests several generations of laboratory evolution. Research often proceeds through unanticipated directions. Reporting such findings responsibly is an essential part of research integrity. Based on a detailed analysis of the genetics of the stocks sampled, we note that our results regarding the chromatin accessibility apply to these lab strains and may not be relevant to the natural populations. Importantly, we want to clarify that the hybrid ancestry was discovered post-hoc after comparison with the natural populations.

To assess chromatin accessibility, ATAC-seq was performed on individuals assumed to be representative of *D. albomicans* and *D. nasuta* parental species. However, the subsequent comparison with whole-genome data from natural populations revealed that these lab-maintained individuals possess significant hybrid ancestry. In particular, both replicates of the ATAC-seq data for *D. albomicans* showed strong signals of *D. nasuta* ancestry, indicating that the identified chromatin accessibility patterns may not accurately represent either species in isolation. Natural populations of *D. nasuta* exhibited tight genetic clustering and limited substructure, consistent with more homogeneous ancestry. In contrast, *D. albomicans* natural populations displayed pronounced substructure and geographical differentiation. As shown by admixture and PCA analyses, laboratory populations were genetically distinct from their wild counterparts, reflecting both hybrid ancestry and potential drift due to bottlenecks and artificial selection under lab conditions. This divergence underscores the importance of validating lab stocks' genetic identity before conducting genome-wide analyses.

The admixture plots and PCA discussed above provide a quantitative description of the hybrid ancestry of the individuals sampled for the ATAC-seq assay. While the experimental design assumed pure parental individuals, subsequent analysis revealed that the samples were genetically admixed hybrids. The two biological replicates differed substantially in their levels of hybrid ancestry. They introduced the potential for genotype-driven differences in observed genomic or transcriptomic patterns unrelated to the experimental treatment or biological condition of interest.

These findings underscore a critical challenge in evolutionary and functional genomics: the assumption of genetic purity in laboratory-maintained strains can obscure or confound biological interpretations. Despite rigorous experimental controls, the biological replicates used for ATAC-seq represent genetically distinct hybrid backgrounds. As a result, any differential chromatin accessibility observed may reflect genetic background effects rather than true species-specific differences. Combining admixture analysis with PCA provides a robust framework for interpreting functional genomic data in the context of complex ancestry. These insights call for greater scrutiny when selecting or maintaining model organisms and highlight the need to integrate population-genetic validation into functional genomics workflows. Ultimately, this study demonstrates that integrating genomic background into experimental design and interpretation is not just ideal but essential.

The choice of reference genome influences the mapping of sequencing reads. Differences in alignment quality and read assignment were observed across parental species references. This bias affects downstream analysis, particularly in admixed or highly divergent genomes. Because reference genome choice and variable ancestry contribute to signal variability, separating biological effects from technical artefacts is challenging. While steps were taken to compare mappings across multiple references, some observed differences may still reflect mapping artefacts rather than underlying biological processes. Therefore, the results presented here are specific to the hybrid individuals analysed and cannot be generalised to either of the presumed parental species. The biological patterns observed must be interpreted in the context of hybrid genomic architecture. Despite these limitations, the study offers valuable insights into the challenges of working with admixed genomes and highlights the need for careful consideration of ancestry and mapping strategies in genomic analyses.

In summary, these extensive ancestry inference analyses establish that the lab stocks of *D. albomicans* (Okinawa strain, Texas collection, USA, 3045.11) and *D. nasuta* (Coorg, India 201.009) collected from the Drosophila Stock Center, University of Mysore, India, appear to be hybrids compared to the natural populations of *D. albomicans* and *D. nasuta* sampled. For consistency, we refer to these as hybrids, as either identifying with *D. albomicans* or *D. nasuta* based on the identity ascribed to them by the stock centre and their hybrid nature identified by our analysis. Therefore, the samples AAT1 and AAT2 are called hybrids identifying with *D. albomicans* despite having mostly *D. nasuta* natural population-like ancestry. Similarly, the samples ANT1 and ANT2 are called hybrids identifying with *D. nasuta* despite having varying levels of *D. albomicans* and *D. nasuta*-like ancestry. Importantly, the lab populations are so different from the natural species that they could be assigned species or subspecies status. Such examples of lab strains being different from natural populations are documented for various organisms and have implications for the study of pathogenesis (Fux et al. 2005; Kumar et al. 2014). Lab strains can diverge enough to lose representativeness, but formal reclassification usually requires an ecological or clinical context (Borneman et al. 2011; Peris et al. 2023). These cases underscore how eukaryotic "lab strains" or industrial isolates—once considered variants of well-known species—can represent cryptic biodiversity. Researchers have formally recognised several new species through genome sequencing and mating tests, particularly within Saccharomyces yeasts.

Classical inbred mouse strains (e.g., C57BL/6, BALB/c, 129) trace their ancestry to wild Mus musculus, incorporating contributions from subspecies domesticus, musculus, castaneus, and even molossinus. These hybrid origins mean different lab strains effectively represent distinct subspecies mosaics, often clustering clearly by subspecies origin (Yang et al. 2011). Genetic analyses confirm that most inbred strains group into clades aligning with wild subspecies: domesticus, musculus, castaneus, or even the outgroup *Mus spretus*, reinforcing that lab strains represent genetically distinct lineages akin to subspecies (Harr 2006). Despite this, nomenclature guidelines deliberately avoid formal subspecies designation for lab mice; instead, they use strain names (e.g., C57BL/6J, PWK/PhJ) registered via institutions like The Jackson Laboratory. Lab adaptations often mimic domestication, with changes in behaviour or development, but do not always come with clear reproductive isolation, a key criterion for subspecies or species designation. Many lab populations originate from hybrids of wild relatives, making taxonomic labelling murky—whether to treat them as new subspecies or just inbred mosaic strains.

**Validation of lab population divergence using independent samples**

All our analysis in the previous section was based on comparing the four ATAC-seq samples and six re-sequenced samples obtained from the Drosophila Stock Center, University of Mysore, for the current project with natural populations. The high levels of sequence divergence between the natural populations and these lab-sampled individuals suggest the build-up of differences over hundreds of generations. Nonetheless, we wanted to validate the prevalence of admixture in the stocks sampled by independent groups to rule out the possibility of mislabelling or mixing of samples during DNA extraction, library preparation and analysis steps. A search of the Short Read Archive revealed that a recent paper (DSouza et al. 2021) reported genome sequencing and assembly for the reference strains and certain cytoraces (**Supplementary Table S2**).

The admixture analysis was performed again, including these six additional samples. The order of the individuals remains the same as before, with the additional six individuals at the right end. Only one individual for each strain and cytorace is available in the public database. Therefore, we had to restrict our analysis to this set of individuals. At a K of 3, the natural populations of *D. albomicans* and *D. nasuta* form distinct clades. Nonetheless, the *D. albomicans* population from India appears admixed with *D. nasuta*. All the lab population members form a distinct clade with differing levels of *D. albomicans* and *D. nasuta* natural population ancestry.

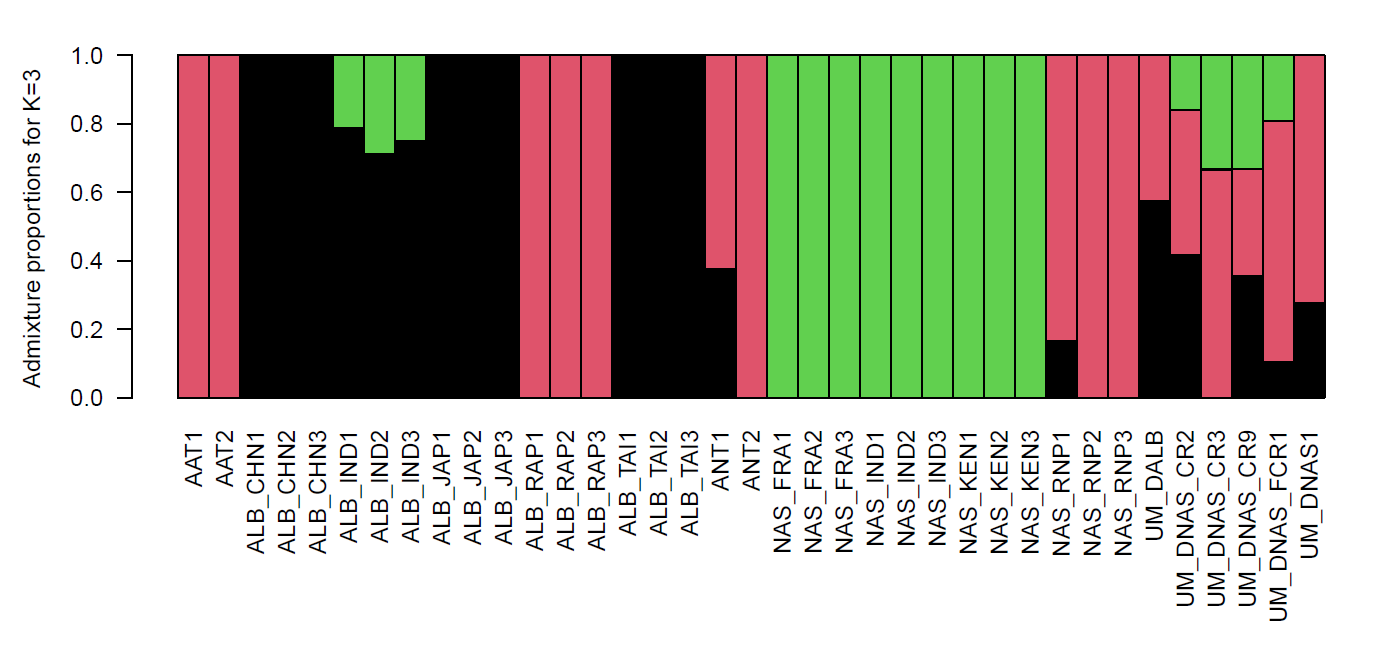

At higher values of K, the mosaic ancestry of the lab populations appears to become evident.

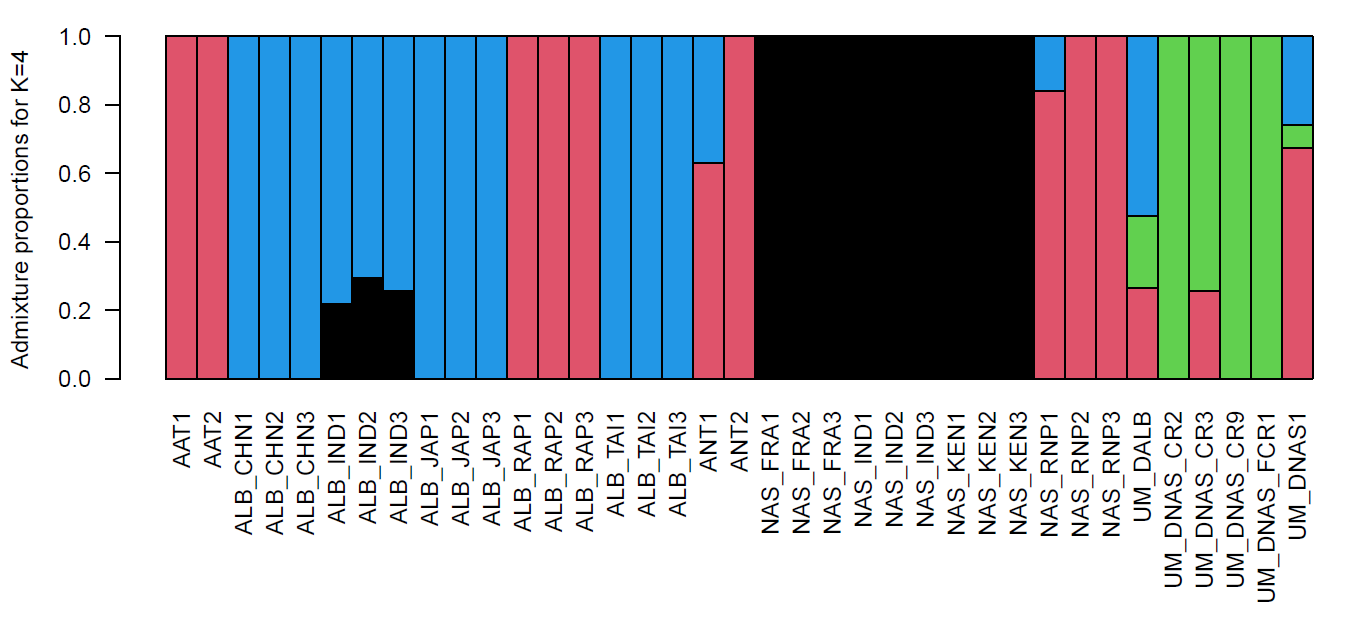

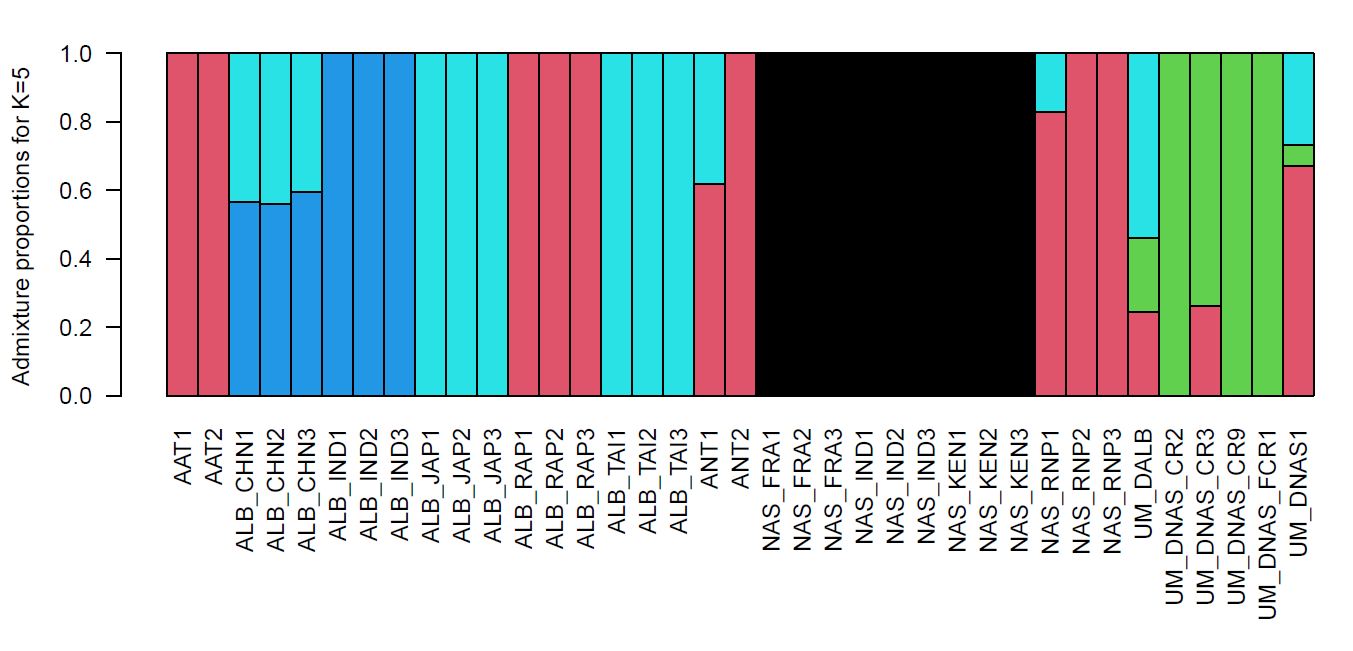

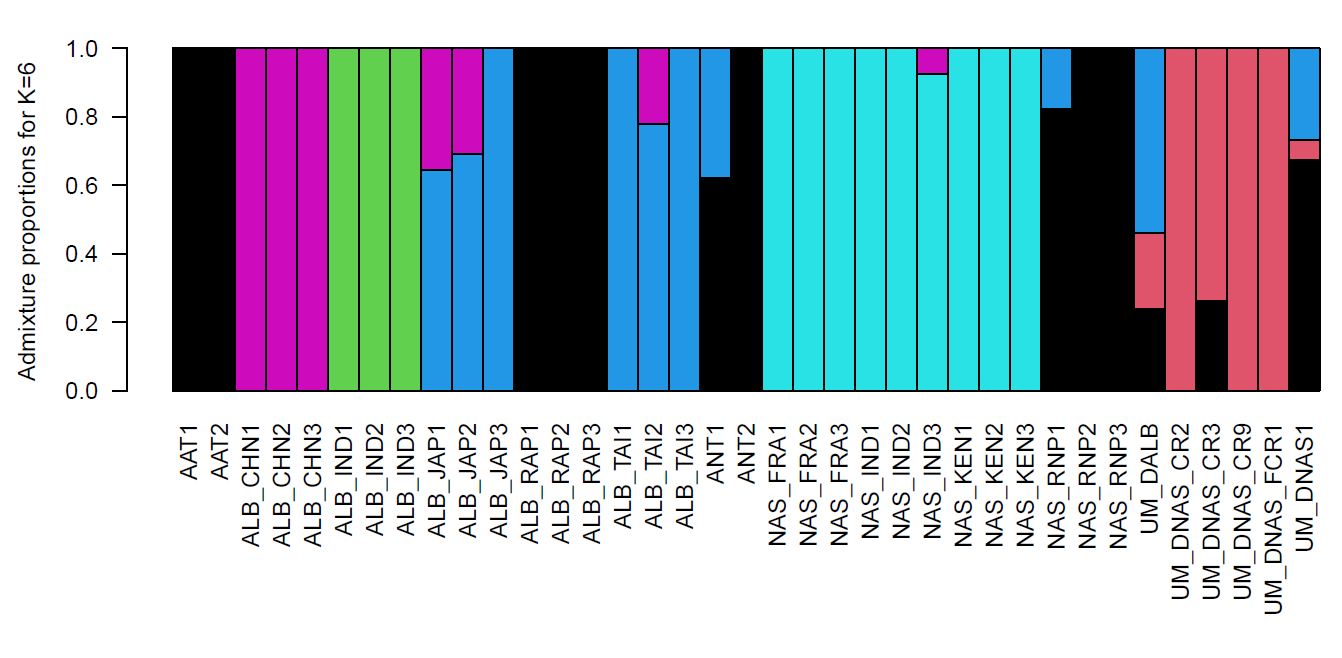

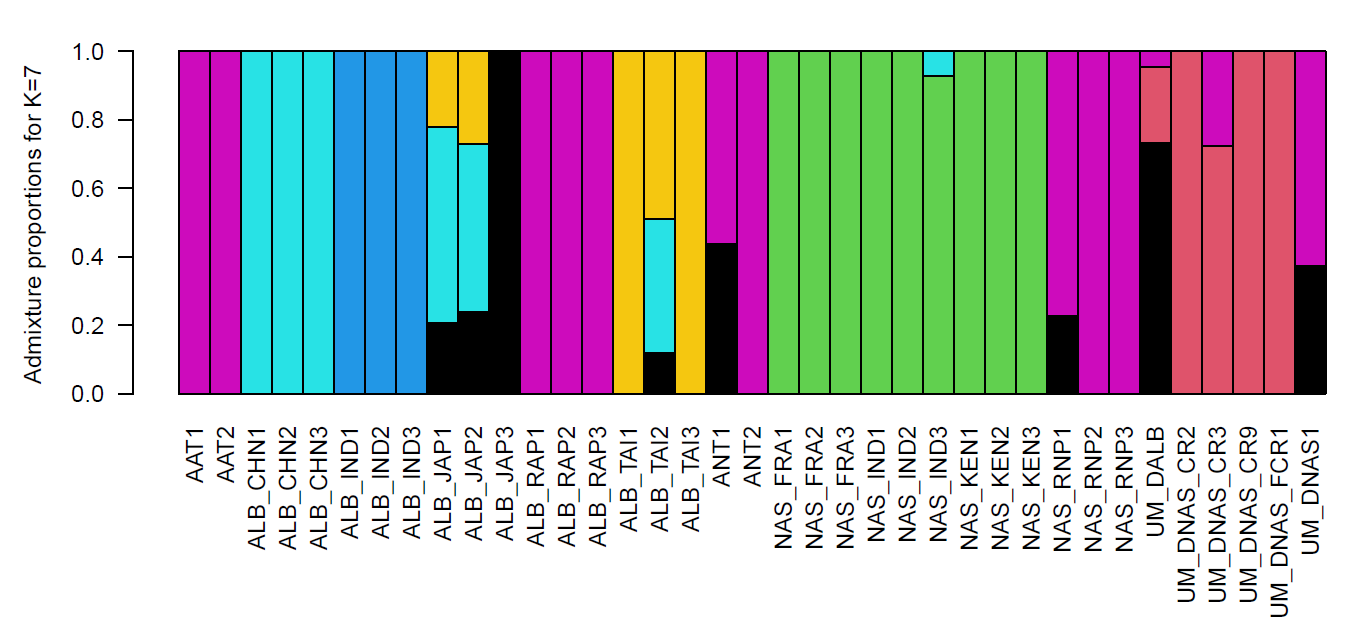

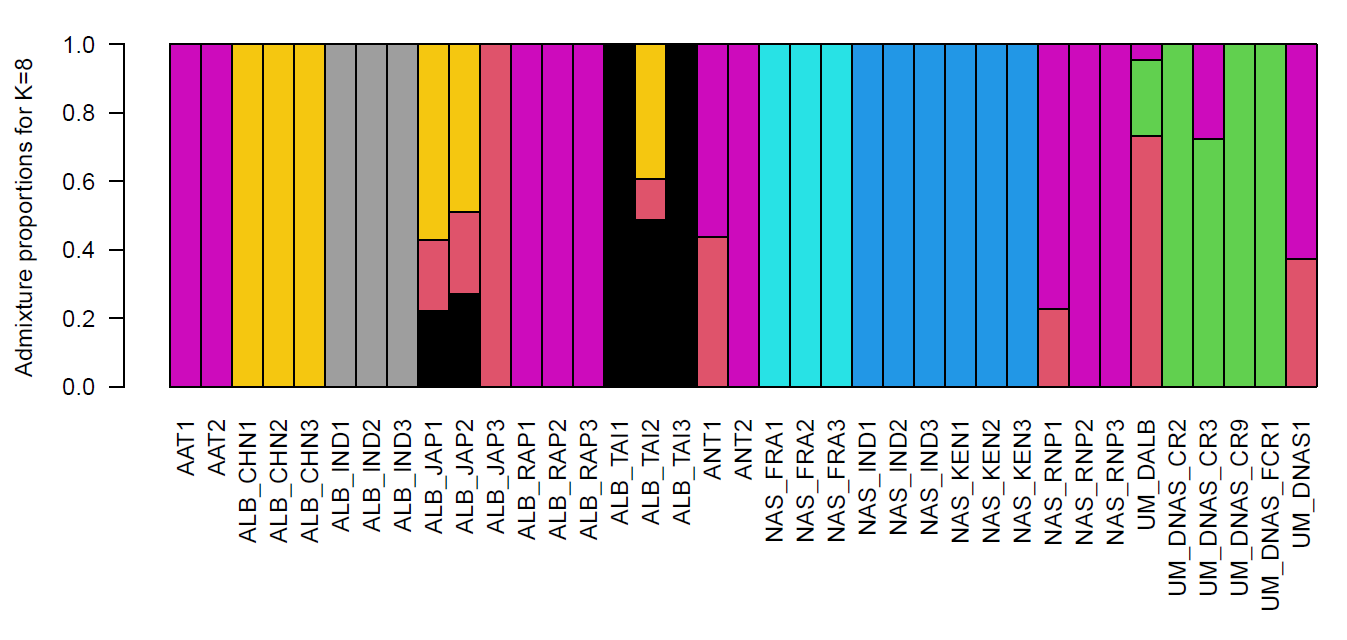

The consistent clustering of the samples sequenced as part of our study and previously published sequences from the University of Mysore allows the validation of a distinct lab population genetically distinct from the natural populations whose genome sequences are reported. We note that additional sampling of other lab strains and more extensive sampling of natural populations will allow a more nuanced understanding of the ancestral origins of the *D. albomicans* and *D. nasuta* stocks maintained at the Drosophila Stock Center, University of Mysore, India.

**References:**

Borneman AR, Desany BA, Riches D, et al (2011) Whole-Genome Comparison Reveals Novel Genetic Elements That Characterize the Genome of Industrial Strains of Saccharomyces cerevisiae. PLoS Genet 7:e1001287. https://doi.org/10.1371/JOURNAL.PGEN.1001287

DSouza S, Ponnanna K, Chokkanna A, Ramachandra N (2021) Illumina short-read sequencing data, de novo assembly and annotations of the Drosophila nasuta nasuta genome. Data Br 34:. https://doi.org/10.1016/j.dib.2020.106674

Fux CA, Shirtliff M, Stoodley P, Costerton JW (2005) Can laboratory reference strains mirror “real-world” pathogenesis? Trends Microbiol 13:58–63. https://doi.org/10.1016/j.tim.2004.11.001

Harr B (2006) Genomic islands of differentiation between house mouse subspecies. Genome Res 16:730. https://doi.org/10.1101/GR.5045006

Korneliussen TS, Albrechtsen A, Nielsen R (2014) ANGSD: Analysis of Next Generation Sequencing Data. BMC Bioinformatics 15:. https://doi.org/10.1186/S12859-014-0356-4

Kumar N, Lad G, Giuntini E, et al (2014) Bacterial genospecies that are not ecologically coherent: population genomics of Rhizobium leguminosarum. Open Biol 5:. https://doi.org/10.1098/RSOB.140133

Peris D, Ubbelohde EJ, Kuang MC, et al (2023) Macroevolutionary diversity of traits and genomes in the model yeast genus Saccharomyces. Nat Commun 14:690. https://doi.org/10.1038/S41467-023-36139-2

Skotte L, Korneliussen TS, Albrechtsen A (2013) Estimating individual admixture proportions from next generation sequencing data. Genetics 195:693–702. https://doi.org/10.1534/GENETICS.113.154138

Yang H, Wang JR, Didion JP, et al (2011) Subspecific origin and haplotype diversity in the laboratory mouse. Nat Genet 43:648–55. https://doi.org/10.1038/ng.847
